# Dissecting the genetic architecture of flowering and maturity time in almond (*Prunus dulcis*): heritability estimates and breeding value predictions from historical data

**DOI:** 10.1101/2025.06.06.658198

**Authors:** MM Gomez-Abajo, Federico Dicenta, Pedro José Martínez-García

**Affiliations:** Fruit Breeding Group. Department of Plant Breeding, Centro de Edafología y Biología Aplicada del Segura-Spanish National Research Council (CEBAS-CSIC). Campus Universitario Espinardo, E-30100 Murcia, Spain

**Keywords:** breeding, almond, flowering, maturity, inheritance, genetic trends, mixed models

## Abstract

Almond (*Prunus dulcis*) is a major nut crop with high genetic complexity due to its heterozygosity and self-incompatibility. In this study, genetic parameters for flowering and maturity time—two key complex traits in almond breeding—were estimated using classical methods (midparent-offspring regression and between/within family variance components) as well as a Bayesian linear mixed model. A comprehensive dataset from the CEBAS-CSIC Almond Breeding Program (CC-ABP), comprising over 17,500 individuals and more than 30 years of historical phenotypic records, was used to generate the first complete pedigree and evaluate trait inheritance. Narrow-sense and broad-sense heritability estimates were obtained, showing substantial variation across traits and methods, with the highest values derived from classical approaches. Bayesian mixed models, implemented via the MCMCglmm R package, allowed for the estimation of breeding values (EBVs), repeatability, and variance components under unbalanced data conditions. Repeatability estimates ranged from 0.15 to 0.56. EBVs were calculated for all individuals, including those without phenotypic records, enabling the construction of trait-specific rankings for early or late flowering and maturity. Reliability values were used to refine these rankings, improving the accuracy of parental selection. Genetic trends based on EBVs revealed changes in breeding objectives over time and highlighted the limited genetic progress achieved through phenotypic selection alone. The integration of the complete pedigree, EBVs, and trait rankings offers a robust framework to optimize crossing strategies. This work lays the groundwork for incorporating genomic selection into future almond breeding efforts, improving selection efficiency for traits of agronomic importance.

## Introduction

The almond (*Prunus dulcis* (Mill.) D.A. Webb), belonging to the subgenus *Amygdalus* within the genus *Prunus* (Rosaceae), is believed to have originated in the Eastern Mediterranean and Southwest Asia, with subsequent expansion into Central Asia and the Western Mediterranean (Pérez de los Cobos et al., 2023). Its cultivation is concentrated in regions with Mediterranean-type climates, with the United States, Australia, and Spain as the main producers. Almonds are classified botanically as drupes; however, unlike other Prunus fruits such as peach or cherry, the economically relevant part is the seed or kernel (Gradziel and Martínez-Gómez, 2013). Almond is predominantly self-incompatible and requires cross-pollination to achieve high yields.

In this sense, self-compatibility has been one of the main goals of almond breeding (Dicenta and García, 1993; Gradziel and Kester, 1998; Ortega and Dicenta, 2003; Socias i Company and Felipe, 1992; Vargas et al., 1998). Since the 20th century, several breeding programs have emerged, notably in Spain (CEBAS-CSIC, IRTA, CITA), the United States (UC Davis), and Australia (University of Adelaide), leading to the release of numerous cultivars such as “Penta”, “Florida” and “Alaska” (CEBAS-CSIC), “Constantí”, “Vairo” and “Marinada” (IRTA), “Belona”, “Soleta” and “Mardía” (CITA), “Sweetheart” and “Winters” (UC Davis) and “Capella”, “Carina” and “Maxima” (Adelaide). A key breeding objective in Spain has been the development of late-flowering cultivars to reduce frost damage (Dicenta et al., 2005; Socias et al., 1999). As a result, flowering time has become a critical trait, influenced by chilling and heat requirements and highly dependent on environmental conditions (Egea et al., 2003; Prudencio et al., 2021). With current climate change, early flowering cultivars are gaining interest for their adaptability to new production areas (Freitas et al., 2023; Alonso Segura et al., 2017).

Other traits of interest include flower density, maturity time, and productivity (Dicenta et al., 1993; Grasselly, 1972; Grasselly and Crossa-Raynaud, 1980; Kester and Asay, 1975; Sánchez-Pérez et al., 2007a; Vargas et al., 1984). Flower density correlates strongly with yield and is a reliable productivity indicator. Early maturity is advantageous in warmer areas to avoid heat stress and in colder areas to reduce the risk of fungal infection and facilitate post-harvest handling (Dicenta et al., 2003, 2016; Connell et al., 2010). These traits are considered complex and polygenic, with high environmental sensitivity. Thus, understanding their genetic architecture is essential to develop efficient breeding strategies. Traditional methods such as mass selection and hybridization have introduced variability (Gradziel and Kester, 1996; Gradziel and Martínez-Gómez, 2013; Socias i Company, 1998), but phenotypic selection alone may be inefficient due to environmental influence and low heritability.

The integration of molecular markers has enabled early selection (marker-assisted selection (MAS)), identification of cultivars, determination of genetic variability (Bouhadida et al., 2011; Dirlewanger et al., 2004; Gradziel and Martínez-Gómez, 2013; Martínez-Gómez et al., 2003; Ortega and Dicenta, 2004; Sánchez-Pérez et al., 2010) and QTL mapping (Fernández i Martí et al., 2013; Más-Gómez et al., in press; Pérez de los Cobos et al., 2024; Sánchez-Pérez et al., 2007b, 2012; Tavassolian et al., 2010), while high-throughput tools like SNP arrays (Duval et al., 2023) and long-read sequencing have enhanced genome resolution and epigenetic analysis (Fresnedo-Ramírez et al., 2023). These advances pave the way for genomic selection, which has already improved efficiency in species like eucalyptus, apple, and apricot (Resende et al., 2012; Kumar et al., 2012; Nsibi et al., 2020).

In the meantime, pedigree-based prediction offers a robust and cost-effective strategy to estimate breeding values (BVs), especially for long-generation crops (Crossa et al., 2006; Oakey et al., 2006). Based on the infinitesimal additive model (Fisher, 1919), this approach allows the calculation of BLUPs to predict genetic merit. The breeding value, defined as twice the deviation of an offspring’s mean phenotype from the population mean under random mating (Falconer, 1983), reflects the additive genetic potential passed to progeny. Its reliability, expressed as the squared correlation between true and estimated BVs, depends on the amount and quality of pedigree and phenotypic data (Gorjanc et al., 2015; Yang and Su, 2016). Higher reliability leads to more accurate selection and greater genetic gain per unit time. No previous studies have applied this approach to almond, although it has proven successful in other species (Martínez-García et al., 2017; Piepho et al., 2008). Additionally, the genetic trend, defined as the change in average BVs over time, is a useful parameter to assess breeding progress and predict future gain (Rutkoski, 2019).

The aim of this study was to estimate breeding values, their reliability, and the genetic parameters of heritability and repeatability for four complex traits (initial, full, and final flowering time, and maturity time) and one categorical trait (flower density) in almond. Repeated measures over multiple years were used to estimate repeatability. Genetic trends were analyzed to assess the evolution of the CC-ABP breeding population.

Rankings were established based on EBVs and on their reliability, to improve selection accuracy.

## Material and Methods

### Plant material

The breeding populations were grown in the experimental field of CEBAS-CSIC in Santomera, Murcia (SE of Spain). It consisted of 17,581 almond trees that constitute CC-ABP.

### Phenotypic Traits

Initial, full and final flowering times were scored in Julian Days, when 5%, 50% and 95% flowers of a tree were open, respectively. Full flowering time is the most common trait used to determine the flowering time of a cultivar. The maturity time was also established in Julian Days when the mesocarp was open in 95% of the fruits on the tree. Flower density was scored visually using an ordinal scale from 0 (no flowers) to 5 (very high density of flowers).

### Pedigree Analysis

For the construction of the pedigree and calculation of the number of founders, the Python program PyPedal (Cole, 2007) was used. Inbreeding coefficients and an ordered pedigree were obtained using the R package pedigreemm (Vazquez et al., 2010).

### Statistical Analysis

A complete descriptive statistical analysis was performed to summarize and to explain the data distribution (e.g., minimum, maximum, average values). This also allowed us to identify phenotypic outliers, eliminating those that could affect the consistency of the data. Narrow-sense heritability (h^2^) was estimated using two traditional methods. The first method was midparent-offspring regression (Falconer, 1983), where the regression coefficient (β_1_) provides a direct estimate of h^2^. The second method involved partitioning variance components between and within families, following the approach described by Kearsey (1965). In this framework, both broad-sense heritability (H^2^ = σ²_F_/σ²_F_+σ²_e_) and narrow-sense heritability (h^2^ = 2σ²_F_/σ²_F_+σ²_e_) were derived, with the distinction lying in the genetic interpretation of the family variance component (σ²_F_). Under the assumption σ²_F_ = ½σ^2^_a_+¼σ^2^_d_, and considering dominance variance (σ^2^_d_) as negligible, σ²_F_ approximates half of the additive genetic variance (σ^2^_a_). Thus, multiplying σ²_F_ by 2 yields an estimate of h^2^. The ratio h^2^ / H^2^ was used to assess the proportion of additive genetic variance relative to total genetic variance, with values close to one indicating a minimal contribution of non-additive effects.

In contrast to traditional methods, a third approach was employed to estimate heritability and to partition genetic variance components for the target traits, following the framework proposed by Martínez-García et al. (2017). For traits following a normal distribution, a linear mixed model was fitted assuming a Gaussian distribution. In contrast, for non-normally distributed traits, such as flower density, a threshold model was used. This model relates the observed categorical data to an underlying multivariate normal distribution via a probit link function (Mrode and Thompson, 2005; Sorensen et al., 1995; Xu and Xu, 2006). In this case, the residual variance was fixed at 1.0, as is standard for categorical traits. Additionally, the repeated phenotypic records typical in almond breeding enabled the inclusion of both genetic effects and permanent environmental effects in the model. This structure accounts for the repeated measurements taken on the same individuals across years. The separation of these effects was possible through the use of the complete pedigree dataset, encompassing 17,581 individuals from the breeding program.

The linear model used was defined as y_ijklm_ = µ + age_i_ + year_j_ + plot_l_ + *a*_kl_ + pe_kl_ + *e*_ijklm_, where *y*_ijklm_ was the m-th observation on the k-th tree, in the l-th plot, during the j-th year, and from the i-th age class. The model includes the overall mean (µ), random effects of age class (age_i_), year (year_j_), and plot (plot_l_), as well as the additive genetic effect (*a*_kl_), the permanent environmental effect (pe_kl_), and the residual error (e_ijklm_). All explanatory variables were treated as random with weakly informative priors due to the large number of levels. The additive genetic effects, *a*_kl_, were assumed to follow a multivariate normal distribution *a*_kl_ ∼ *N*(0, *A*σ^2^_a_), where *A* is the numerator relationship matrix for 17,581 trees. Permanent environmental effects, pe_kl_ ∼ *N*(0, *I*σ^2^_pe_), and residuals e_ijklm_ ∼ *N*(0, *I*σ^2^_e_). Age, year, and plot were modeled as independent random effects with variances σ²_year_, σ²_age_, and σ²_plot_, respectively. The main goal was to estimate μ, predict the random effects, and calculate the variances σ²_a_, σ²_pe_ and σ²_e_. These estimates allowed for the calculation of narrow-sense heritability (h^2^ = σ²_a_/(σ²_a_+σ²_pe_+σ²_e_)) and repeatability (r = (σ² +σ²) / (σ² +σ² +σ²)). Notably, variances attributed to non-genetic factors, such as year, plot, and age, were excluded from the denominator when calculating heritability and repeatability. Consequently, the estimates of these genetic parameters were considered within specific year, plot, and age groups Bayesian inference was performed using the MCMCglmm package in R (Hadfield, 2015), following the methodology of Martínez-García et al. (2017). Fixed effects were assigned diffuse normal priors, and random effects followed multivariate normal distributions. Variance components were estimated using inverse-Wishart priors. Each MCMC chain ran for 500,000 iterations with a 50,000 burn-in and thinning interval of 200, yielding 2,250 posterior samples. Model convergence was assessed using trace plots, autocorrelation diagnostics, and posterior density inspection (Plummer et al., 2006). The high posterior density (HPD) intervals provided measures of precision for the estimates. Narrower HPD intervals indicated higher confidence in the posterior mode estimates. Genetic trends for EBVs were evaluated through linear regression over years, and Pearson’s correlations were calculated among traits. Inbreeding coefficients were estimated using the R package pedigreemm.

## Results

### Data Overview

Historical phenotypic data from the CC-ABP breeding population, covering over 30 years of evaluation, were used to analyze the selected traits. The dataset included individuals with varying numbers of annual records, from 1,713 individuals evaluated in a single year (e.g., selection “D00_002”) to long-term cultivars such as “Marcona,” “Ferragnès,” and “Desmayo Largueta,” which had records spanning over 20 years. The cultivar with the most extensive data was “R1000,” with 26 observations across 17 years. On average, each individual had 2.45 records, with 1,711 individuals represented by a single observation (e.g., “D00_010”). The average age of individuals in the breeding population was 3.73 years, with “Achaak” and “Desmayo Largueta” being the oldest at 33 years.

Regarding the traits studied, the number of families, individuals, and years varied across traits (Table 1). Flower density and full flowering time were the most consistently recorded, each with 30 years of data. Flower density had the widest representation, with 8,088 individuals and 344 families. For full flowering time, maturity time, and flower density, the family with the most data was “R1000 × Desmayo Largueta”. In contrast, for initial and final flowering time, the most extensively recorded family was “Ramillete × Mono,” while “Ramillete × Tuono” had the highest number of individual trees observed (Table 2). Trait distribution is shown in Figure 1. All traits displayed an approximately normal distribution, except for flower density, which is an ordinal categorical trait. For the breeding population (individuals with data from at least two years), the average Julian day values were: 50 for initial flowering, 57 for full flowering, 60 for final flowering, and 221 for maturity time. The average flower density score was 2.02.

**Fig. 1.**
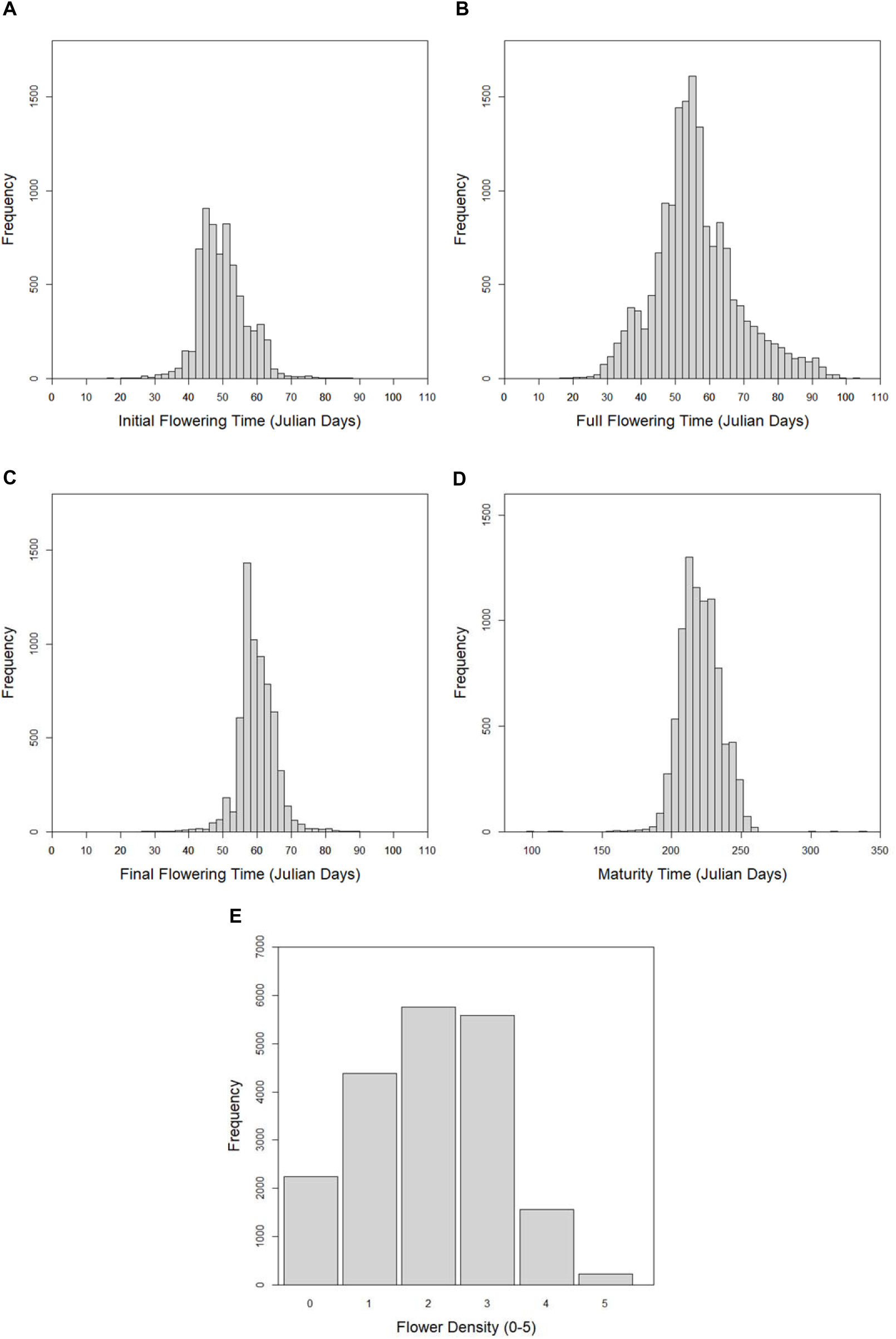
Frequency histograms of initial **(A)**, full (**B)** and final **(C)** flowering times, and maturity time **(D)** in Julian days. **E.** Frequency histogram of flower density scored from 0 (no flowers) to 5 (very high density).

**Table 1.** Number of total data, number of individuals and families with data and number of years of study for each trait.

|  | Total data | Individuals | Families | Years |
| --- | --- | --- | --- | --- |
| Initial flowering time | 6,554 | 2,636 | 77 | 10 |
| Full flowering time | 16,381 | 7,382 | 338 | 30 |
| Final flowering time | 6,520 | 2,616 | 73 | 9 |
| Maturity time | 8,524 | 4,974 | 306 | 28 |
| Flower density | 19,729 | 8,088 | 344 | 30 |

**Table 2.** Top five families with the highest number of data recorded, and the number of individuals, for each trait.

| Family | Data | Individuals |
| --- | --- | --- |
| **Initial flowering time** |  |  |
| Ramillete x Mono | 302 | 109 |
| Ramillete x Tuono | 282 | 125 |
| Tuono x Ferragnès | 269 | 116 |
| Atocha x Genco | 226 | 93 |
| Primorskii x Garrigues | 220 | 78 |
| **Full flowering time** |  |  |
| R1000 x Desmayo Langueta | 982 | 304 |
| Desmayo Langueta x R1000 | 607 | 209 |
| Ramillete x Mono | 302 | 109 |
| Tuono x Ferragnès | 283 | 116 |
| Ramillete x Tuono | 282 | 125 |
| **Final flowering time** |  |  |
| Ramillete x Mono | 302 | 109 |
| Ramillete x Tuono | 282 | 125 |
| Tuono x Ferragnès | 268 | 116 |
| Atocha x Genco | 226 | 93 |
| Primorskii x Garrigues | 220 | 78 |
| **Maturity time** |  |  |
| R1000 x Desmayo Langueta | 422 | 228 |
| Ramillete x Tuono | 207 | 124 |
| Ramillete x Mono | 199 | 108 |
| S5133 x Antoñeta | 178 | 74 |
| Primorskii x Garrigues | 147 | 75 |
| **Flower density** |  |  |
| R1000 x Desmayo Langueta | 2,007 | 616 |
| Desmayo_Langueta x R1000 | 617 | 209 |
| Ramillete x Tuono | 375 | 125 |
| Tuono x Ferragnès | 361 | 116 |
| Ramillete x Mono | 327 | 109 |

Among cultivars and selections, “Desmayo Largueta” and “Marcona” had the most extensive records for flowering traits, with up to 23 years of data for full flowering time. For maturity time, the advanced selection “D00_078” stood out with 15 years of data. In terms of phenological extremes, “Desmayo Largueta” showed the earliest average initial flowering (23 Julian days), while “Tardona” was the latest (82 Julian days). For full flowering, “D10_006_B2” was the earliest (25 Julian days) and “D09_305” the latest (93 Julian days). For final flowering, “Desmayo Largueta” again flowered earliest (37 Julian days), and “R1000” latest (85 Julian days). In maturity time, “D07_158” was the earliest individual with an average of 163 Julian days, while “D04_341” was the latest, at 276 Julian days. Regarding flower density, score 2 was the most frequent across the population. A total of 223 individuals achieved the maximum score of 5, while 2,236 scored 0 at least once. The highest average flower density was observed in “D01_631” (4.75), whereas 97 individuals maintained an average score of 0.

### Correlations between traits

In general, the correlations among flowering time traits were very high. The strongest correlation was found between initial and full flowering time (r = 0.96), while the weakest—although still high—was between initial and final flowering time (r = 0.89). In contrast, flower density showed the lowest correlations with the rest of the traits, with most coefficients close to zero. The highest correlation involving flower density was with full flowering time (r = 0.10), which is still considered very low. Maturity time presented weak correlations with the flowering traits. The highest correlations were observed with initial flowering time (r = 0.19) and final flowering time (r = 0.22), while its correlations with full flowering time (r = -0.07) and flower density (r = -0.02) were negative and negligible. All correlation coefficients were statistically significant. Most were extremely significant (*p* < 0.001), except for the correlation between flower density and initial flowering time, which was highly significant (*p* < 0.01), and the correlation between flower density and maturity time, which was significant (*p* < 0.05) (Figure 2).

**Fig. 2.**
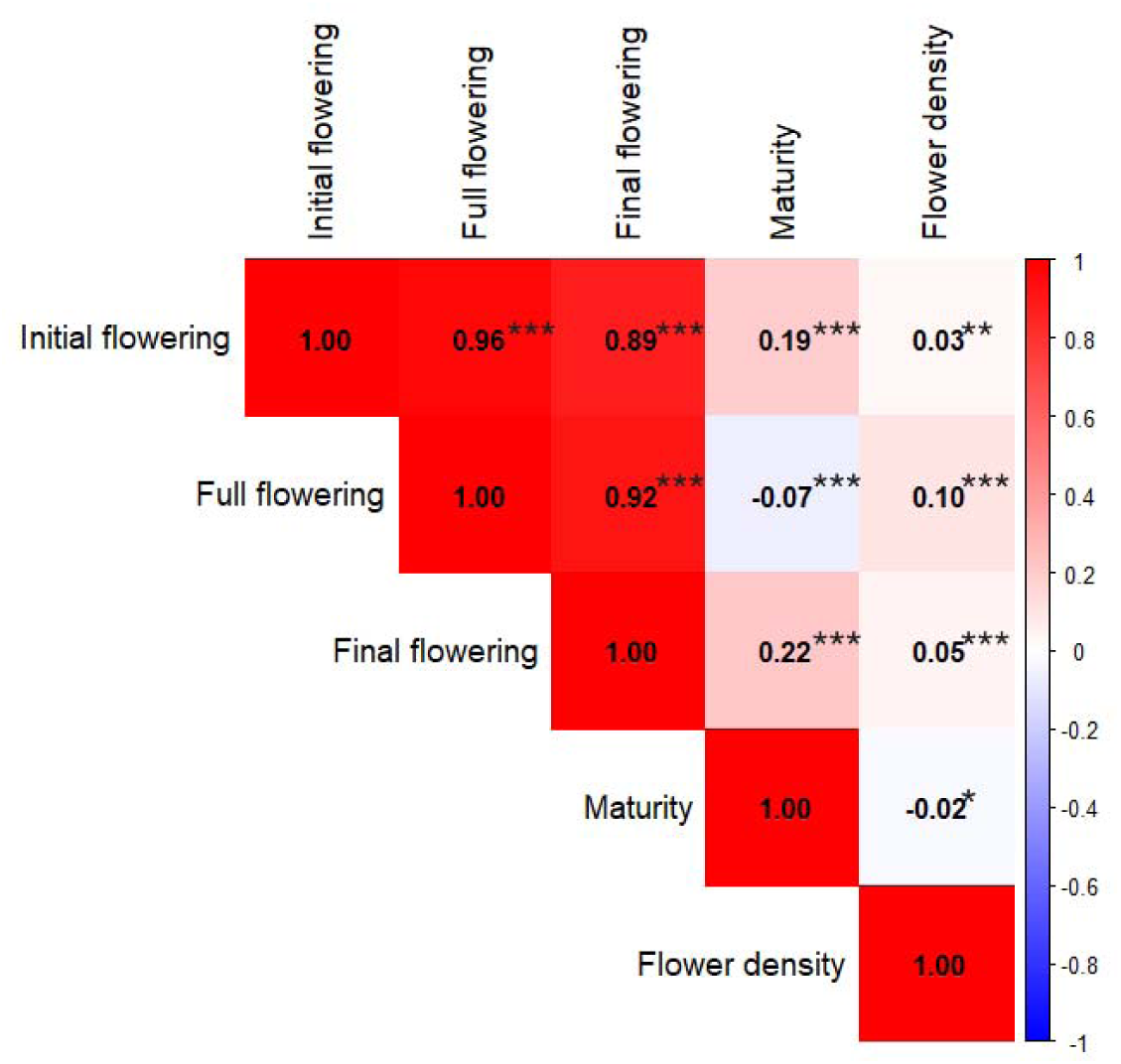
Pearson correlation matrix between all the studied traits by corrplot analysis. Asterisks indicate significance levels: p < 0.05 (*), p < 0.01 (**), and p < 0.001 (***).

### Pedigree Analysis

For this study, the first complete pedigree of the CC-ABP was created, presenting a total of 17,581 individuals. From this set, 17,529 were descendants obtained from a wide range of crosses, including the seven almond cultivars of the CEBAS-CSIC “Antoñeta”, “Marta”, “Penta”, “Makako”, “Tardona”, “Alaska” and “Florida”. The pedigree of these varieties is illustrated in Figure 3. The remaining 52 progenitors were traditional cultivars (with different origins), relative species of almond (such as *Prunus webbii*, *Prunus scoparia* or *Prunus orientalis*) and Spanish ecotypes (such as “ITAP-1”). This set of individuals has been used as parents at least one time in the CC-ABP. After the obtaining of the pedigree, and according to the number of descendants (and relatives) of each individual, the results showed a reduced number of 38 individuals which could be considered as the “main” founders of the CC-ABP. The main contributor to this breeding program is the cultivar “Tuono”, having the largest number of descendants, with a total of 13,899. This pedigree was composed of 432 families, with 36 families having more than 100 descendants. There were 420 full-sib families and 12 half-sib families (with one unknown parent). For the full sib families, “R1000” x “Desmayo Largueta” was the largest family with 633 descendants. Among the half-sib families, the largest family, with 80 descendants, had the cultivar “Tardona” as mother tree. As a result of the cross between “Tuono” and “Ferragnès”, “Antoñeta” and “Marta” were obtained. “Tuono” was also the parent of “Lauranne” and “R1000”. “Lauranne” progeny included “Penta” and “Makako”, while “R1000” was the parent of “Tardona” (Figure 3).

**Fig 3.**
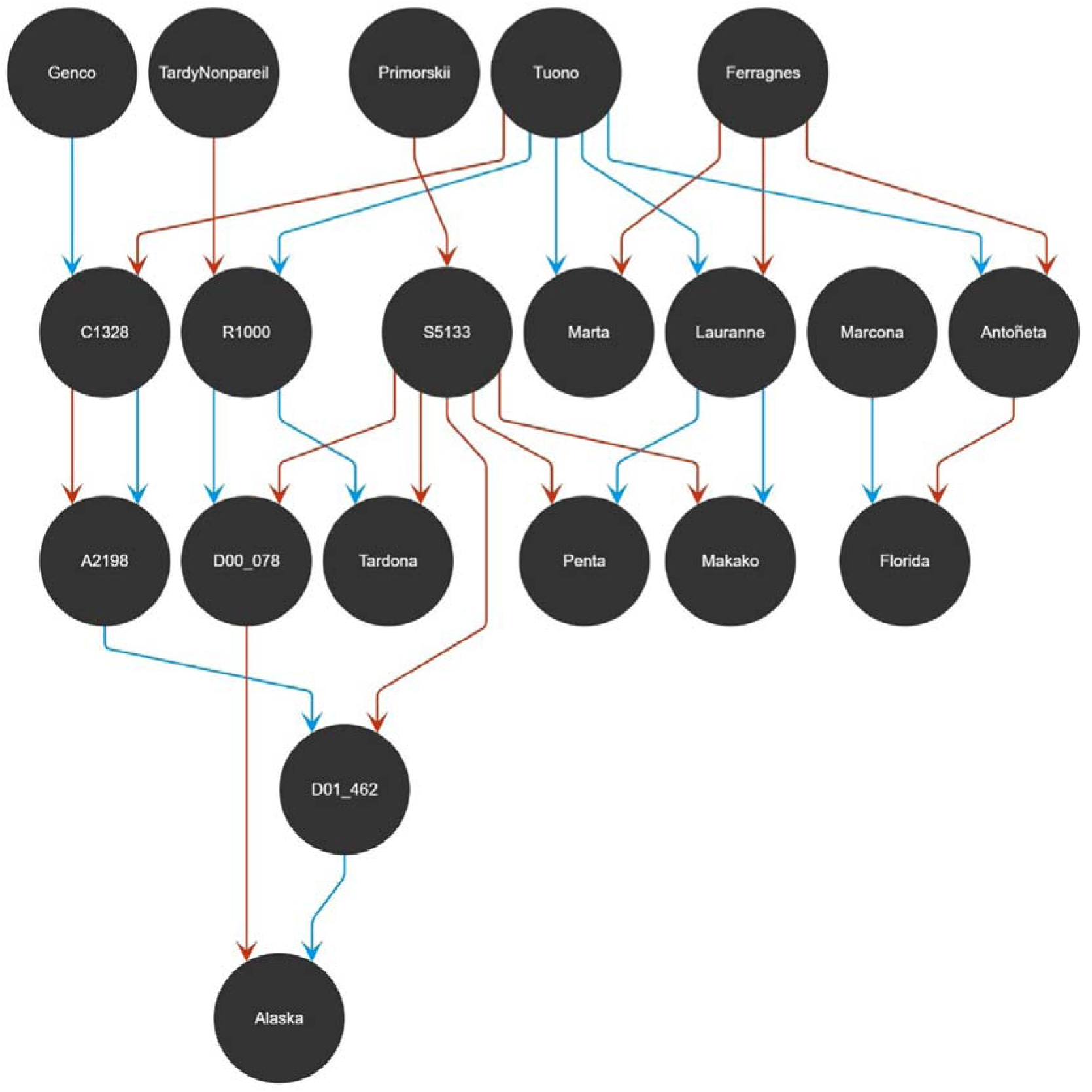
Pedigree diagram of the CEBAS-CSIC cultivars "Antoñeta", "Marta", "Penta", "Makako", "Tardona", “Alaska” and “Florida”. Arrows represent parental contributions: red for female parents (dams) and blue for male parents (sires).

The inbreeding coefficient of the breeding population was low, with an average of 0.10 and 68% of the individuals with an inbreeding coefficient below average. The range of coefficients was from 0 to 0.75 and the highest value was presented by four families obtained by self-pollination of selections “A1180”, “A1340”, “A1342” and “A1353”.

### Estimation of Heritability and genetic parameters

Heritability estimates obtained from both classical and mixed models are summarized in Table 3. The highest narrow-sense heritability values were derived using the between/within family variance components method, with estimates ranging from 0.38 to 1.56. Estimates based on midparent-offspring regression ranged from 0.34 to 1.07. Broad-sense heritability values ranged from 0.19 to 0.77, and were consistently lower than the corresponding narrow-sense estimates. Within the Bayesian linear mixed model framework, maturity time showed the highest narrow-sense heritability (0.42), while final flowering time showed the lowest (0.14).

**Table 3.** Narrow- and broad-sense heritability estimates from classical methods and Bayesian mixed models.

| Trait | Method | $h^2$ | $H^2$ |
| --- | --- | --- | --- |
| **Initial flowering time** | Midparent-offspring regression | 1.07 | - |
|  | Between/within family variance components | 1.41 | 0.71 |
|  | Bayesian mixed model | 0.24 | - |
| **Full flowering time** | Midparent-offspring regression | 0.92 | - |
|  | Between/within family variance components | 1.33 | 0.77 |
|  | Bayesian mixed model | 0.32 | - |
| **Final flowering time** | Midparent-offspring regression | 0.80 | - |
|  | Between/within family variance components | 1.56 | 0.76 |
|  | Bayesian mixed model | 0.14 | - |
| **Maturity time** | Midparent-offspring regression | 0.45 | - |
|  | Between/within family variance components | 1.22 | 0.61 |
|  | Bayesian mixed model | 0.42 | - |
| **Flower density** | Midparent-offspring regression | 0.34 | - |
|  | Between/within family variance components | 0.38 | 0.19 |
|  | Bayesian mixed model | 0.16 | - |

All genetic parameter estimates, including heritability and repeatability for the four traits evaluated, are shown in Table 4. Posterior density and trace plots were examined, revealing symmetrical and non-aberrant curves, confirming the unimodality of posterior densities (Figure S1-S10). The highest repeatability estimate was observed for maturity time (0.56), while the lowest was for final flowering time (0.15). For flower density, the 95% highest posterior density (HPD) interval for heritability ranged from 0.12 to 0.24, indicating high estimation precision. In contrast, initial flowering time showed the widest HPD interval (0.10 to 0.39), reflecting lower accuracy. For repeatability, flower density again presented the narrowest HPD interval (0.20 to 0.34). In comparison, the HPD range for full flowering time and final flowering time was approximately 0.20, and for initial flowering time and maturity time, around 0.30. The permanent environmental variance component showed the most precise estimates across traits, with narrow HPD intervals. For maturity time, the HPD interval for this component was the widest (11.21). Among flowering time traits, the year variance displayed the largest HPD intervals. For maturity time and flower density, the least precise estimates corresponded to plot variance, with intervals ranging from 5.31 to 103.83 and from 0.30 to 2.45, respectively. In terms of the percentage of total variance explained, the year effect was the dominant source for all flowering time traits, ranging from 40.30% to 46.28%. For maturity time, the largest proportion of variance was explained by breeding values (45.12%). In the case of flower density, the largest contributor was the residual variance (23.25%), followed by age (20.70%) and breeding values (19.77%). The permanent environmental effect accounted for between 2.63% (final flowering time) and 15.80% (maturity time) of the total variance.

**Table 4.** Summary of the posterior modes of variance components, heritability, repeatability, and the corresponding 95% highest posterior density (HPD) intervals for the evaluated traits. The proportion of variance explained by each effect is indicated in parentheses.

|  | Posterior mode | HPD 2.5 | HPD 97.5 |
| --- | --- | --- | --- |
| **Initial flowering time** |  |  |  |
| Heritability | 0.24 | 0.10 | 0.39 |
| Repeatability | 0.27 | 0.11 | 0.45 |
| Breeding value variance | 18.85 (35.40%) | 16.42 | 20.76 |
| Permanent effect variance | 3.37 (6.33%) | 2.35 | 4.22 |
| Year variance | 24.65 (46.28%) | 14.06 | 129.24 |
| Age variance | 5.27 (9.90%) | 1.58 | 22.56 |
| Plot variance | 0.11 (0.20%) | 0.00 | 4.11 |
| Residual variance | 1.00 (1.88%) | 1.00 | 1.00 |
| **Full flowering time** |  |  |  |
| Heritability | 0.32 | 0.23 | 0.39 |
| Repeatability | 0.39 | 0.29 | 0.48 |
| Breeding value variance | 36.88 (35.38%) | 32.78 | 39.52 |
| Permanent effect variance | 8.17 (7.84%) | 6.54 | 9.75 |
| Year variance | 42.01 (40.30%) | 24.55 | 76.64 |
| Age variance | 12.87 (12.35%) | 8.29 | 24.23 |
| Plot variance | 3.30 (3.16%) | 1.44 | 15.12 |
| Residual variance | 1.00 (0.96%) | 1.00 | 1.00 |
| **Final flowering time** |  |  |  |
| Heritability | 0.14 | 0.04 | 0.22 |
| Repeatability | 0.15 | 0.04 | 0.25 |
| Breeding value variance | 14.97 (19.10%) | 13.05 | 16.63 |
| Permanent effect variance | 2.06 (2.63%) | 1.24 | 2.76 |
| Year variance | 35.67 (45.50%) | 15.92 | 209.04 |
| Age variance | 23.99 (30.61%) | 8.73 | 97.84 |
| Plot variance | 0.70 (0.89%) | 0.00 | 12.03 |
| Residual variance | 1.00 (1.28%) | 1.00 | 1.00 |
| **Maturity time** |  |  |  |
| Heritability | 0.42 | 0.27 | 0.49 |
| Repeatability | 0.56 | 0.38 | 0.65 |
| Breeding value variance | 77.31 (45.12%) | 66.00 | 88.29 |
| Permanent effect variance | 27.08 (15.80%) | 20.98 | 32.19 |
| Year variance | 22.74 (13.28%) | 13.79 | 50.19 |
| Age variance | 21.02 (12.27%) | 12.49 | 39.97 |
| Plot variance | 22.17 (12.94%) | 5.31 | 103.83 |
| Residual variance | 1.00 (1.28%) | 1.00 | 1.00 |
| **Flower density** |  |  |  |
| Heritability | 0.16 | 0.12 | 0.24 |
| Repeatability | 0.28 | 0.20 | 0.34 |
| Breeding value variance | 0.85 (19.77%) | 0.70 | 1.05 |
| Permanent effect variance | 0.44 (10.23%) | 0.37 | 0.57 |
| Year variance | 0.37 (8.60%) | 0.22 | 0.73 |
| Age variance | 0.89 (20.70%) | 0.45 | 1.61 |
| Plot variance | 0.75 (17.44%) | 0.30 | 2.45 |
| Residual variance | 1.00 (23.25%) | 1.00 | 1.00 |

### Breeding Values and Reliability

Estimated breeding values (EBVs) and their associated reliability were calculated for each individual in the CC-ABP pedigree (Supplemental File 2). For individuals with at least one phenotypic record for the studied traits, EBVs were estimated using both phenotypic data and genetic relationships derived from the pedigree. For the remaining 9,252 individuals without phenotypic observations, EBVs were inferred solely from pedigree information. A ranking of the top 20 genotypes with the highest EBVs for each trait was established (Table 5). This includes the 20 earliest and latest flowering individuals—relevant for breeding in warmer regions with low chilling requirements, and in colder regions where late flowering helps avoid frost damage. The ranking also highlights individuals with earlier or later maturity times, as well as those with the highest EBVs for flower density. Among all individuals in the pedigree, “Desmayo Largueta” had the earliest EBVs for all flowering traits: –21.84 for initial, –26.54 for full, and –17.79 for final flowering time. “D99_646” also showed early flowering for initial (–17.60) and final (–13.09) flowering time. For full flowering, the earliest EBV was observed in “D04_408” (–25.25). The latest flowering genotypes were “D08_382” for initial (31.74), “D05_145” for full (38.95), and “D98_672” for final flowering time (25.50). Regarding maturity, “D20_409” had the earliest EBV (–38.47), and “D10_349” the latest (64.26). The highest EBV for flower density was recorded in “D98_694” (2.82).

**Table 5.**
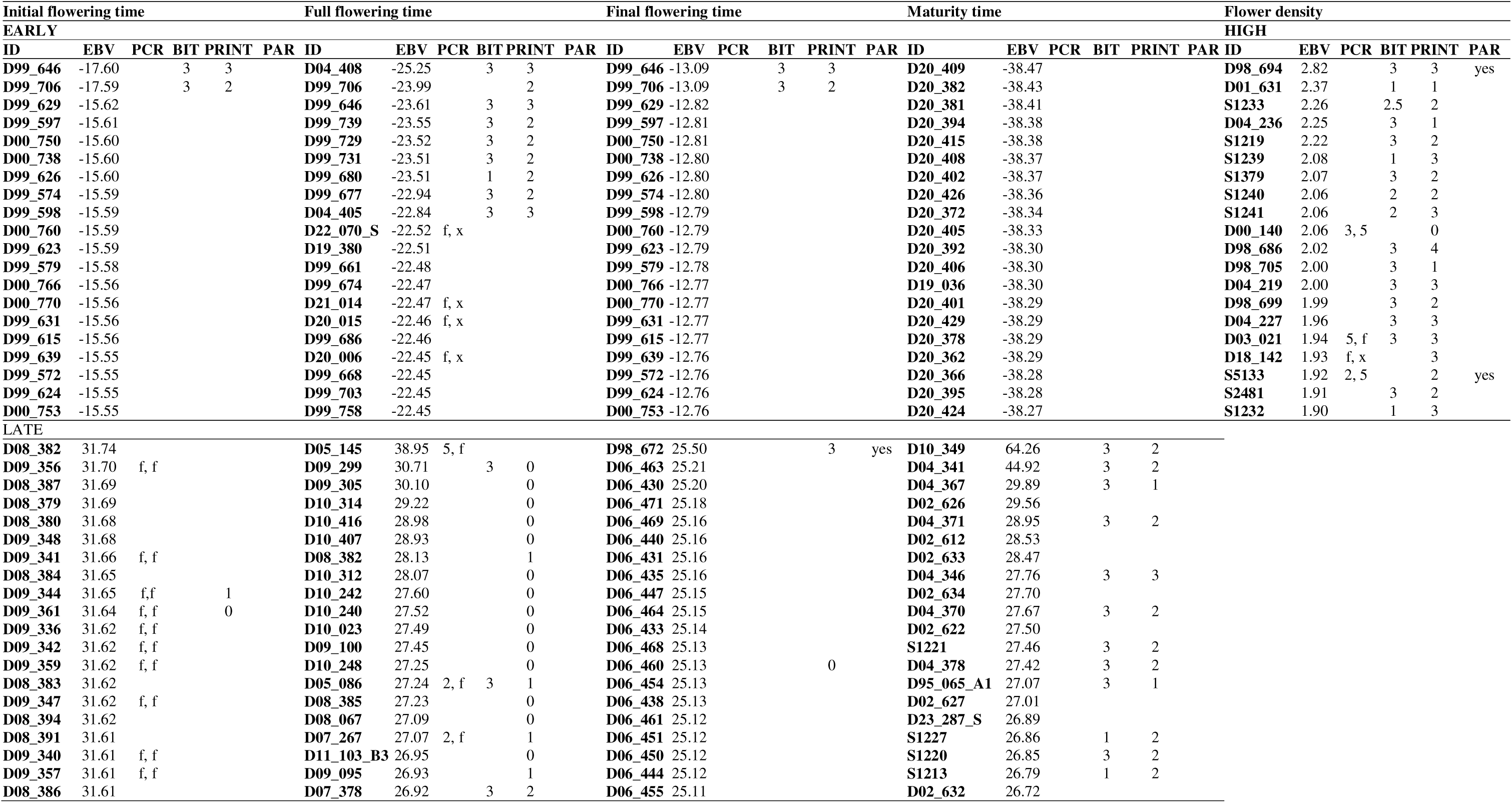
Ranking of the 20 best genotypes of the CC-ABP according to their EBV for each trait, differentiated by early and late and high flower density, with available data on self-compatibility alleles (PCR), where individuals carrying the Sf (f) allele are self-compatible, bitterness (BIT) and production intensity (PRINT), both expressed as the average of all recorded years for each individual, and use as parents (PAR)

Reliability values of EBVs ranged from 0 to values close to 0.90 across all traits. “R1000” showed the highest individual EBV reliability (0.98) for full flowering time. This trait had the highest average reliability (0.62), followed by maturity time (0.52). Final flowering time showed the lowest average reliability (0.37). For each trait, a final ranking was developed based on both EBV and reliability. A threshold reliability of ≥0.8 was applied for full flowering time, and ≥0.5 for the remaining traits. Based on these criteria, 1,196 individuals were identified as promising candidates for advancing full flowering and maturity time (i.e., EBV < 0). For example, “D00_714” had an EBV of –21.98 for full flowering time, and “D17_235” had an EBV of –32.10 for maturity time. A similar filtering strategy was used to select individuals with high flower density, yielding 286 candidates. Among them, selection “S2481” showed the highest EBV for flower density (1.91) (Supplemental File 3). Pearson correlations were calculated between the average EBVs of each trait. All correlations were highly significant (*p* < 0.001). Strong positive correlations (r > 0.90) were observed among the three flowering time traits. Maturity time showed a moderate correlation with flower density (r = 0.30) and a lower correlation with final flowering time (r = 0.14). Correlations between flower density and the flowering time traits ranged from 0.25 to 0.30. Regression analyses revealed a significant temporal trend in EBVs for initial flowering time (*p* = 0.027, R² = 0.17) and flower density (*p* = 0.015, R² = 0.20). No significant trends were detected for full flowering time (*p* = 0.06), final flowering time (*p* = 0.080), or maturity time (*p* = 0.049), all with low coefficients of determination (R² between 0.11 and 0.14) (Figure 4).

**Fig. 4.**
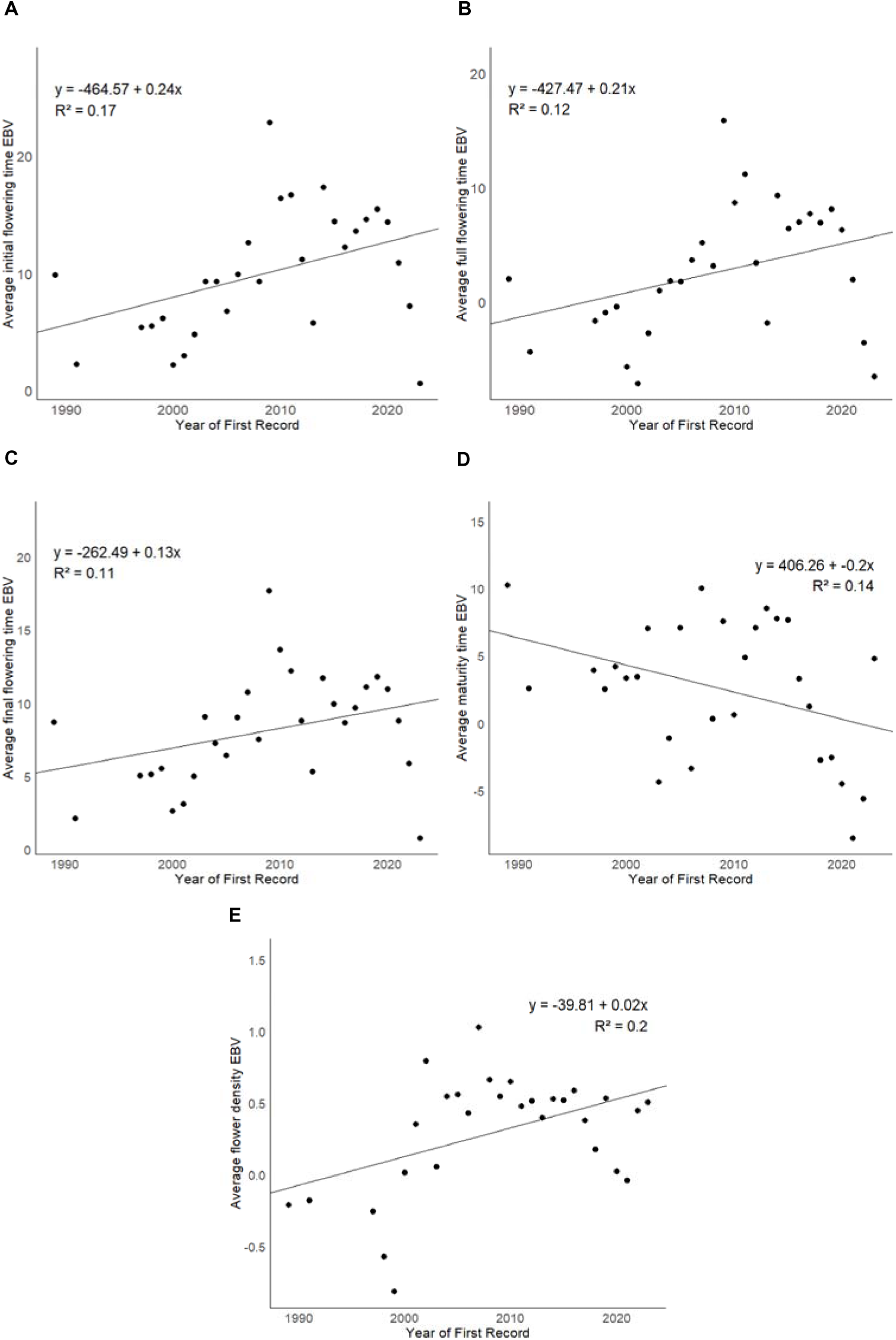
Genetic trend of average estimated breeding values (EBV) for each trait by year of first record**. A.** Genetic trend of EBVs of start of flowering time. **B.** Genetic trend of EBVs of full flowering time. **C.** Genetic trend of EBVs of end of flowering time. **D.** Genetic trend of EBVs of start of maturity time. **E.** Genetic trend of EBVs of flower density.

## Discussion

In this study, a well-defined generalized linear mixed model was developed to estimate breeding values for key traits in almond breeding. This was achieved using the first complete pedigree of the CC-ABP and over 30 years of historical phenotypic records. The extensive amount of phenotypic and pedigree data generated through traditional breeding provided a strong foundation for the construction of robust genetic models. As a result, narrow-sense heritability estimates were obtained for the first time in the CC-ABP using a Bayesian linear mixed model. Rankings of individuals with the best breeding values were also produced. These results will help increase selection efficiency for economically important traits such as flowering and maturity time.

Previous studies have reported heritability estimates and described inheritance patterns for flowering time, maturity time, and flower density in almond (*Prunus dulcis*) using traditional methods (Grasselly, 1972; Kester, 1965; Vargas and Romero, 1988; Dicenta et al., 1993; Dicenta and García, 1993). These studies generally indicated moderate to high heritability for these traits. In the present work, estimates obtained through classical approaches—midparent–offspring regression and between/within family variance partitioning—were largely consistent with those earlier findings, particularly for flowering-related traits. However, it is essential to interpret heritability estimates in the context of their underlying assumptions and limitations. Heritability, whether in the broad sense (H^2^) or narrow sense (h^2^), is a population-specific and environment-specific parameter; it does not represent an intrinsic property of a trait (Falconer and Mackay, 1996). High heritability indicates that, within the studied population and conditions, a large proportion of phenotypic variance is attributable to genetic differences—not that the trait is genetically “determined” or unaffected by the environment. This distinction is often misunderstood, leading to overinterpretation of heritability values (Visscher et al., 2008). Moreover, classical methods like the between/within family approach can overestimate h^2^ in the presence of non-additive genetic effects (dominance, epistasis) or shared environmental effects within families. The assumption that genetic effects are purely additive rarely holds for complex traits, especially in perennial species where long generation times and clonal propagation allow accumulation of epistatic interactions. Cases where narrow-sense heritability estimates exceed broad-sense values are a red flag for model misspecification or confounding effects. In contrast, linear mixed models provide a more flexible framework for estimating heritability, particularly under unbalanced experimental designs with missing data or unequal family sizes—as is common in perennial breeding populations like the CC-ABP. These models can partition variance more accurately and, when extended to genomic data, allow estimation of marker-based heritability using genomic relationship matrices (Lynch and Walsh, 1998). Still, even in such models, failure to account for genotype-by-environment interactions (G×E) or environmental covariates can result in inflated additive variance components.

In other *Prunus* species mixed models have been used (de Souza et al., 1998a, 1998b; Dirlewanger et al., 2012; Calle et al., 2020; Hernández Mora et al., 2017). These studies reinforce the notion that while heritability estimates remain a cornerstone of quantitative genetics, their magnitude reflects both genetic architecture and experimental context. Without comprehensive modeling of genetic and environmental variance—including non-additive and G×E components—heritability estimates may misrepresent the true potential for selection response. In *Prunus avium*, Piaskowski et al. (2018) used genomic-enabled mixed models to estimate heritability for several industry-relevant traits, reporting broad-sense heritability (H²) values of 0.83 for maturity date, 0.77 for fruit firmness, and 0.76 for fruit weight. Notably, these traits are known to be highly influenced by environmental conditions, yet the high heritability values observed highlight the strength of the genetic signal captured in the studied population. The authors also found that non-additive components played a substantial role: epistatic variance accounted for more than 40% of the total genetic variance for maturity date, firmness, and disease response, while dominance variance contributed 34% and 27% of the genetic variance for fruit weight and size, respectively. These results emphasize the importance of using comprehensive genetic models that account for non-additive variance, particularly in clonally propagated perennial species where such effects are more likely to accumulate. Furthermore, they demonstrate that high heritability estimates in complex traits must be interpreted with care, as they may reflect both genuine genetic determinism and methodological factors such as model structure, population composition, and environmental homogeneity.

Trait correlations found in this study were largely consistent with those previously reported. High correlations among flowering traits, and weak to moderate correlations between flower density and maturity traits, align with prior work by Dicenta and García (1992). Interestingly, a weak negative correlation between full flowering and maturity time was observed in this study. This contrasts with previous reports of no correlation (Grasselly, 1972; Sánchez-Pérez et al., 2007a; Sorkheh et al., 2010), as well as with older studies that described positive correlations in almond (Dicenta and García, 1992) and in other species such as walnut and sweet cherry (Hansche et al., 1966, 1972). Historically, a negative correlation between flowering time and flower density was proposed (Grasselly, 1972; Kester, 1965). This assumption persisted until the work of Grasselly and Olivier (1985), who used “Tardy Nonpareil” in breeding crosses and found no such correlation. In the present study, a significant but very low positive correlation was observed between flowering time traits and flower density. While the high correlation among some traits would allow for the application of multi-trait BLUP models (Piepho et al., 2008), this approach was not implemented here due to computational limitations, model convergence issues, and insufficient data for some traits (Bauer and Léon, 2008). Moreover, the use of models developed for continuous traits to predict categorical traits such as flower density may reduce predictive accuracy (Kizilkaya et al., 2014).

This study highlights the value of leveraging historical phenotypic records in perennial crop breeding, especially when combined with a corrected pedigree (Marrano et al., 2019). Pedigree-based analysis, long used in animal and plant breeding (Henderson, 1984; Crossa et al., 2006; Burgueño et al., 2007), enables the monitoring of genetic relationships and inbreeding levels. The current analysis of the CC-ABP showed a higher mean inbreeding coefficient than in previous pedigree studies in almond (Pérez de los Cobos et al., 2021), mainly due to the repeated use of a few founder genotypes (Falconer, 1983), such as “Tuono”. Although pedigree-based methods have been widely applied in other tree crops such as walnut (Martínez-García et al., 2017), peach (de Souza et al., 2000; Fresnedo-Ramírez et al., 2016), apple (Iwanami et al., 2008; Kouassi et al., 2009), blueberry (Cellon et al., 2018) or mango (Hardner et al., 2012), this is the first known report applying this methodology to almond. The EBV-based rankings generated in this study revealed a significant gap between genetic merit and actual parental use. For instance, high-EBV individuals were rarely used in past crosses. Traditional cultivars like ‘Ferragnès’ and ‘R1000’—often selected for late flowering— ranked relatively low in breeding value, highlighting a mismatch between selection goals and actual genetic potential.

When applying reliability thresholds, only a small proportion of the population met selection criteria for early flowering, early maturity, and high flower density simultaneously. This emphasizes the need to improve parental selection strategies in future breeding. One of the main limitations of this study is the lack of multi-environment data to properly estimate genotype-by-environment (G × E) interactions (Lynch and Walsh, 1998). Proper evaluation of G × E effects requires replicated genotypes across diverse environments, as demonstrated in peach (Cirilli et al., 2020), apple (Jung et al., 2020), and other crops using clonal trials (Ward et al., 2019; Chacón et al., 2024 Ferrão et al., 2021; Gezan et al., 2016; Nelson et al., 2018; Poupon et al., 2023). In almond, these kinds of populations are not available, something that should be considered in the future by the almond breeding community.

Genetic trends over time reflect the historical evolution of the CC-ABP breeding objectives. Despite six generations of selection (1985–2024), most traits showed weak or non-significant genetic trends, highlighting the limited genetic gain achieved through traditional phenotypic selection. A shift toward late flowering cultivars occurred around 2000 to reduce frost risk, followed by a more recent focus on early flowering to meet lower chilling requirements. Maturity time was the only trait with a clear negative trend, aligning with a growing preference for earlier maturing cultivars. The lack of improvement in flower density may result from its categorical nature, environmental influence, and negative association with late flowering. Overall, these findings underscore the slow progress of conventional breeding in woody species.

In future perspectives, the integration of genomic tools such as the Axiom™ 60K SNP array for almond (Duval et al., 2023), combined with high-throughput phenotyping technologies, will allow the incorporation of molecular information to refine pedigree data and construct genomic relationship matrices. This will facilitate the calculation of genomic estimated breeding values (GEBVs), enabling the capture of Mendelian sampling variation and increasing the accuracy of selection. The CC-ABP includes a wide genetic diversity in flowering and maturity times—from very early to extra-late cultivars—and continues to develop crosses targeting key objectives, including self-compatibility, early maturity, adaptation to climate change, and drought resistance through rootstock development. These resources and breeding efforts provide a strong foundation for implementing genomic selection and advancing molecular breeding strategies in almond.

## Conclusions

This study highlights the advantages of using linear mixed model framework with Bayesian inference, to effectively partition genetic variance components in populations with unbalanced data such as the CC-ABP. These relatively low heritability values reflect the complexity of the genetic architecture of the studied traits. This may help to explain the difficulty associated with breeding based only on phenotypic data. The selection scheme should be optimized by selecting the best parents to maximize genetic gain. Breeding values are an essential tool to enhance both the accuracy and effectiveness of parental selection in breeding programs. Moreover, reliable pedigree data are crucial for robust estimation of genetic parameters and breeding values. The incorporation of genomic data will improve the accuracy of breeding values and help correct pedigree information, facilitating the implementation of genomic selection in almond.

## Acknowledgments

M.M.G.-A. acknowledges the grant PRE2022-105038 funded by MICIU/AEI/10.13039/501100011033 and by “ESF Investing in your future”.

## Author contributions

P.J.M-G. and F.D. conceived and designed the research. M.M.G.-A. performed the research and analysed the data. P.J.M.-G., M.M.G.-A. and F.D. wrote the manuscript. All authors read and approved the manuscript.

## References

1. Alonso Segura, J. M., Socias i Company R., and Kodad, O. (2017). Late-blooming in almond: A controversial objective. Scientia Horticulturae, 224, 61–67. 10.1016/j.scienta.2017.05.036

2. Bauer, A. M., and Léon, J. (2008). Multiple-trait breeding values for parental selection in self-pollinating crops. Theoretical and Applied Genetics, 116(2), 235–242. 10.1007/s00122-007-0662-6

3. Bermann, M., Aguilar, I., Lourenco, D., Misztal, I., and Legarra, A. (2023). Reliabilities of estimated breeding values in models with metafounders. Genetics Selection Evolution, 55(1), 6. 10.1186/s12711-023-00778-2

4. Bouhadida, M., Moreno, M. Á., Gonzalo, M. J., Alonso, J. M., and Gogorcena, Y. (2011). Genetic variability of introduced and local Spanish peach cultivars determined by SSR markers. Tree Genetics & Genomes, 7(2), 257–270. 10.1007/s11295-010-0329-3

5. Burgueño, J., Crossa, J., Cornelius, P. L., Trethowan, R., McLaren, G., and Krishnamachari, A. (2007). Modeling Additive × Environment and Additive × Additive × Environment Using Genetic Covariances of Relatives of Wheat Genotypes. Crop Science, 47(1), 311–320. 10.2135/cropsci2006.09.0564

6. Calle, A., Cai, L., Iezzoni, A., and Wünsch, A. (2020). Genetic Dissection of Bloom Time in Low Chilling Sweet Cherry (Prunus avium L.) Using a Multi-Family QTL Approach. Frontiers in Plant Science, 10. 10.3389/fpls.2019.01647

7. Cellon, C., Amadeu, R. R., Olmstead, J. W., Mattia, M. R., Ferrao, L. F. V., and Munoz, P. R. (2018). Estimation of genetic parameters and prediction of breeding values in an autotetraploid blueberry breeding population with extensive pedigree data. Euphytica, 214(5), 87. 10.1007/s10681-018-2165-8

8. Chacón, J. G., Fernandez, G. E., and Isik, F. (2024). Genetic Variation in Yield and Fruit Weight Among Strawberry (Fragaria ×ananassa) Cultivars and the Interaction With Year and Location Effect. Plant Breeding, n/a(n/a). 10.1111/pbr.13246

9. Cirilli, M., Micali, S., Aranzana, M. J., Arús, P., Babini, A., Barreneche, T., Bink, M., Cantin, C. M., Ciacciulli, A., Cos-Terrer, J. E., Drogoudi, P., Eduardo, I., Foschi, S., Giovannini, D., Guerra, W., Liverani, A., Pacheco, I., Pascal, T., Quilot-Turion, B., Verde, I., Rossini, L., and Bassi, D. (2020). The Multisite PeachRefPop Collection: A True Cultural Heritage and International Scientific Tool for Fruit Trees. Plant Physiology, 184(2), 632–646. 10.1104/pp.19.01412

10. Clark, S. A., and van der Werf, J. (2013). Genomic Best Linear Unbiased Prediction (gBLUP) for the Estimation of Genomic Breeding Values. En C. Gondro, J. van der Werf, & B. Hayes (Eds.), Genome-Wide Association Studies and Genomic Prediction (pp. 321–330). Humana Press. 10.1007/978-1-62703-447-0_13

11. Cole, J. B. (2007). PyPedal: A computer program for pedigree analysis. Computers and Electronics in Agriculture, 57(1), 107–113. 10.1016/j.compag.2007.02.002

12. Connell, J. H., Gradziel, T. M., Lampinen, B. D., Micke, W. C., and Floyd, J. (2010). Harvest maturity of almond cultivars in California’s Sacramento Valley. 94.

13. Crossa, J., Burgueño, J., Cornelius, P. L., McLaren, G., Trethowan, R., and Krishnamachari, A. (2006). Modeling Genotype × Environment Interaction Using Additive Genetic Covariances of Relatives for Predicting Breeding Values of Wheat Genotypes. Crop Science, 46(4), 1722–1733. 10.2135/cropsci2005.11-0427

14. Crossa, J., Campos, G. de los, Pérez, P., Gianola, D., Burgueño, J., Araus, J. L., Makumbi, D., Singh, R. P., Dreisigacker, S., Yan, J., Arief, V., Banziger, M., and Braun, H.-J. (2010). Prediction of Genetic Values of Quantitative Traits in Plant Breeding Using Pedigree and Molecular Markers. Genetics, 186(2), 713–724. 10.1534/genetics.110.118521

15. de Souza, V. A., Byrne, D. H., and Taylor, J. F. (1998a). Heritability, genetic and phenotypic correlations, and predicted selection response of quantitative traits in peach: I. An analysis of several reproductive traits. Journal of the American Society for Horticultural Science, 123(4), 598–603.

16. de Souza, V. A., Byrne, D. H., and Taylor, J. F. (1998b). Heritability, genetic and phenotypic correlations, and predicted selection response of quantitative traits in peach: II. An analysis of several fruit traits. Journal of the American Society for Horticultural Science, 123(4), 604–611.

17. de Souza, V. A., Byrne, D. H., and Taylor, J. F. (2000). Predicted breeding values for nine plant and fruit characteristics of 28 peach genotypes. Journal of the American Society for Horticultural Science, 125(4), 460–465.

18. Dicenta, F., Egea, J., Ortega, E., Martínez-Gómez, P., Sánchez-Pérez, R., Rubio, M., Martínez-García, P. J., Gómez, E. M., del Cueto, J., Sánchez-Prudencio, A., and Cremades, T. (2016). Almond Breeding: Important issues and challenges for reserach. *Options Méditerranéennes*, Série A, 119, 23–28.

19. Dicenta, F., and García, J. E. (1992). Phenotypical correlations among some traits in almond. Journal of Genetics & Breeding, 46(3), 241–246.

20. Dicenta, F., and García, J. E. (1993). Inheritance of self-compatibility in almond. Heredity, 70(3), 313–317. 10.1038/hdy.1993.45

21. Dicenta, F., Garcia, J. E., and Carbonell, E. A. (1993). Heritability of flowering, productivity and maturity in almond. Journal of Horticultural Science, 68(1), 113–120. 10.1080/00221589.1993.11516334

22. Dicenta, F., García-Gusano, M., Ortega, E., and Martinez-Gómez, P. (2005). The possibilities of early selection of late-flowering almonds as a function of seed germination or leafing time of seedlings. Plant Breeding, 124(3), 305–309. 10.1111/j.1439-0523.2005.01090.x

23. Dicenta, F., Martinez-Gómez, P., Martínez-Pato, E., and Gradziel, T. M. (2003). Screening for Aspergillus flavus resistance in almond. HortScience, 38(2), 266–268.

24. Dirlewanger, E., Graziano, E., Joobeur, T., Garriga-Calderé, F., Cosson, P., Howad, W., and Arús, P. (2004). Comparative mapping and marker-assisted selection in Rosaceae fruit crops. Proceedings of the National Academy of Sciences, 101(26), 9891–9896. 10.1073/pnas.0307937101

25. Dirlewanger, E., Quero-García, J., Le Dantec, L., Lambert, P., Ruiz, D., Dondini, L., Illa, E., Quilot-Turion, B., Audergon, J.-M., Tartarini, S., Letourmy, P., and Arús, P. (2012). Comparison of the genetic determinism of two key phenological traits, flowering and maturity dates, in three Prunus species: Peach, apricot and sweet cherry. Heredity, 109(5), 280–292. 10.1038/hdy.2012.38

26. Durel, C. E., Laurens, F., Fouillet, A., and Lespinasse, Y. (1998). Utilization of pedigree information to estimate genetic parameters from large unbalanced data sets in apple. Theoretical and Applied Genetics, 96(8), 1077–1085. 10.1007/s001220050842

27. Duval, H., Coindre, E., Ramos-Onsins, S. E., Alexiou, K. G., Rubio-Cabetas, M. J., Martínez-García, P. J., Wirthensohn, M., Dhingra, A., Samarina, A., and Arús, P. (2023).

28. Development and Evaluation of an AxiomTM 60K SNP Array for Almond (Prunus dulcis). Plants, 12(2), Article 2. 10.3390/plants12020242

29. Egea, J., Ortega, E., Martı nez-Gómez, P., and Dicenta, F. (2003). Chilling and heat requirements of almond cultivars for flowering. Environmental and Experimental Botany, 50(1), 79–85. 10.1016/S0098-8472(03)00002-9

30. Falconer, D.S. (1983). Introduction to Quantitative Genetics. Oliver and Boyd Ltd.

31. Falconer, D.S., & Mackay, T.F.C. (1996). Introduction to Quantitative Genetics (4th ed.). Longman.

32. Fernández i Martí, A., Font i Forcada, C., and Socias i Company, R. (2013). Genetic analysis for physical nut traits in almond. Tree Genetics & Genomes, 9(2), 455–465. 10.1007/s11295-012-0566-8

33. Ferrão, L. F. V., Amadeu, R. R., Benevenuto, J., de Bem Oliveira, I., and Munoz, P. R. (2021). Genomic Selection in an Outcrossing Autotetraploid Fruit Crop: Lessons From Blueberry Breeding. Frontiers in Plant Science, 12. 10.3389/fpls.2021.676326

34. Fisher, R. A. (1919). The Correlation between Relatives on the Supposition of Mendelian Inheritance. Earth and Environmental Science Transactions of The Royal Society of Edinburgh, 52(2), 399–433. 10.1017/S0080456800012163

35. Freitas, T. R., Santos, J. A., Silva, A. P., and Fraga, H. (2023). Reviewing the Adverse Climate Change Impacts and Adaptation Measures on Almond Trees (Prunus dulcis). Agriculture, 13(7), Article 7. 10.3390/agriculture13071423

36. Fresnedo-Ramírez, J., Anderson, E. S., D’Amico-Willman, K., and Gradziel, T. M. (2023). A review of plant epigenetics through the lens of almond. The Plant Genome, 16(4), e20367. 10.1002/tpg2.20367

37. Fresnedo-Ramírez, J., Frett, T. J., Sandefur, P. J., Salgado-Rojas, A., Clark, J. R., Gasic, K., Peace, C. P., Anderson, N., Hartmann, T. P., and Byrne, D. H. (2016). QTL mapping and breeding value estimation through pedigree-based analysis of fruit size and weight in four diverse peach breeding programs. Tree Genetics & Genomes, 12(2), 25.

38. Gezan, S. A., de Carvalho, M. P., and Sherrill, J. (2016). Statistical methods to explore genotype-by-environment interaction for loblolly pine clonal trials. Tree Genetics & Genomes, 13(1), 1. 10.1007/s11295-016-1081-0

39. Gianola, D., and Fernando, R. L. (1986). Bayesian Methods in Animal Breeding Theory. Journal of Animal Science, 63(1), 217–244. 10.2527/jas1986.631217x

40. Goddard, M. e., and Hayes, B. j. (2007). Genomic selection. Journal of Animal Breeding and Genetics, 124(6), 323–330. 10.1111/j.1439-0388.2007.00702.x

41. Gorjanc, G., Bijma, P., and Hickey, J. M. (2015). Reliability of pedigree-based and genomic evaluations in selected populations. Genetics Selection Evolution, 47(1), 65. 10.1186/s12711-015-0145-1

42. Gradziel, T. M., and Kester, D. E. (1996). Genetic improvements. In Almond Production Manual (pp. 70–75).

43. Gradziel, T. M., and Kester, D. E. (1998). BREEDING FOR SELF-FERTILITY IN CALIFORNIA ALMOND CULTIVARS. Acta Horticulturae, 470, 109–117. 10.17660/ActaHortic.1998.470.15

44. Gradziel, T. M., and Martínez-Gómez, P. (2013). Almond Breeding. In Plant breeding reviews (Vol. 37, pp. 207–258).

45. Grasselly, C. (1972). L’amandier: Caractères morphologiques et physiologiques des variétés, modalité de leurs transmissions chez les hybrides de première génération. University of Bordeaux.

46. Grasselly, C., and Crossa-Raynaud, P. (1980). L’amandier. G.P. Maisonneuve et Larose.

47. Grasselly, C., and Olivier, G. (1985). Avancement du programme de creation de varietes d’amandier autocompatible. Options Méditerranéennes. Série Etudes, 2.

48. Hadfield, J. (2015). MCMCglmm: MCMC generalised linear mixed models. Royal Society Open Science, 3(160), 087.

49. Hansche, P. E., Beres, W., and Brooks, R. M. (1966). Heritability and genetic correlation in the sweet cherry. Journal of the American Society for Horticultural Science, 88, 173–183.

50. Hansche, P. E., Beres, W., and Forde, H. I. (1972). Estimates of quantitative genetic properties of walnut and their implications for cultivar improvement. Journal of the American Society for Horticultural Science, 97(2), 279–285.

51. Hardner, C. M., Bally, I. S. E., and Wright, C. L. (2012). Prediction of breeding values for average fruit weight in mango using a multivariate individual mixed model. Euphytica, 186(2), 463–477. 10.1007/s10681-012-0639-7

52. Henderson, C. R. (1984). Applications of linear models in animal breeding (Vol. 462).

53. Hernández Mora, J. R., Micheletti, D., Bink, M., Van de Weg, E., Cantín, C., Nazzicari, N., Caprera, A., Dettori, M. T., Micali, S., Banchi, E., Campoy, J. A., Dirlewanger, E., Lambert, P., Pascal, T., Troggio, M., Bassi, D., Rossini, L., Verde, I., Quilot-Turion, B., Laurens. F., Arús, P. and Aranzana, M. J. (2017). Integrated QTL detection for key breeding traits in multiple peach progenies. BMC Genomics, 18(1), 404. 10.1186/s12864-017-3783-6

54. Iwanami, H., Moriya, S., Kotoda, N., Takahashi, S., and Abe, K. (2008). Estimations of Heritability and Breeding Value for Postharvest Fruit Softening in Apple. 10.21273/JASHS.133.1.92

55. Jung, M., Roth, M., Aranzana, M. J., Auwerkerken, A., Bink, M., Denancé, C., Dujak, C., Durel, C.-E., Font i Forcada, C., Cantin, C. M., Guerra, W., Howard, N. P., Keller, B., Lewandowski, M., Ordidge, M., Rymenants, M., Sanin, N., Studer, B., Zurawicz, E., Laurens, F., Patocchi, A., Muranty, H. (2020). The apple REFPOP—a reference population for genomics-assisted breeding in apple. Horticulture Research, 7, 189. 10.1038/s41438-020-00408-8

56. Kearsey, M. J. (1965). Biometrical analysis of a random mating population: A comparison of five experimental designs. Heredity, 20(205–235)

57. Kester, D. E. (1965). Inheritance of time of bloom in certain progenies of almond. Proceedings, 87, 214–221.

58. Kester, D. E., and Asay, R. (1975). Almonds. In Advances in Fruit Breeding (pp. 367–384). Purdue University Press.

59. Kizilkaya, K., Fernando, R. L., and Garrick, D. J. (2014). Reduction in accuracy of genomic prediction for ordered categorical data compared to continuous observations. Genetics Selection Evolution, 46(1), 37. 10.1186/1297-9686-46-37

60. Kouassi, A. B., Durel, C.-E., Costa, F., Tartarini, S., van de Weg, E., Evans, K., Fernandez-Fernandez, F., Govan, C., Boudichevskaja, A., Dunemann, F., Antofie, A., Lateur, M., Stankiewicz-Kosyl, M., Soska, A., Tomala, K., Lewandowski, M., Rutkovski, K., Zurawicz, E., Guerra, W., and Laurens, F. (2009). Estimation of genetic parameters and prediction of breeding values for apple fruit-quality traits using pedigreed plant material in Europe. Tree Genetics & Genomes, 5(4), 659–672. 10.1007/s11295-009-0217-x

61. Kumar, S., Bink, M. C. A. M., Volz, R. K., Bus, V. G. M., and Chagné, D. (2012). Towards genomic selection in apple (Malus ×domestica Borkh.) breeding programmes: Prospects, challenges and strategies. Tree Genetics & Genomes, 8(1), 1–14. 10.1007/s11295-011-0425-z

62. Lynch, M., and Walsh, B. (1998). Genetics and analysis of quantitative traits (Vol. 1).

63. Marrano, A., Martínez-García, P. J., Bianco, L., Sideli, G. M., Di Pierro, E. A., Leslie, C. A., Stevens, K. A., Crepeau, M. W., Troggio, M., Langley, C. H., and Neale, D. B. (2019). A new genomic tool for walnut (Juglans regia L.): Development and validation of the high-density Axiom^TM^ J. regia 700K SNP genotyping array. Plant Biotechnology Journal, 17(6), 1027–1036. 10.1111/pbi.13034

64. Martínez-García, P. J., Famula, R. A., Leslie, C., McGranahan, G. H., Famula, T. R., and Neale, D. B. (2017). Predicting breeding values and genetic components using generalized linear mixed models for categorical and continuous traits in walnut (Juglans regia). Tree Genetics & Genomes, 13(5), 109. 10.1007/s11295-017-1187-z

65. Martínez-Gómez, P., Arulsekar, S., Potter, D., and Gradziel, T. M. (2003). An extended interspecific gene pool available to peach and almond breeding as characterized using simple sequence repeat (SSR) markers. Euphytica, 131(3), 313–322. 10.1023/A:1024028518263

66. Más-Gómez, J., Gómez-López, F. J., Rubio, M., Gómez-Abajo, M. M., Dicenta, F., and Martínez-García, P. J. (in press). Integration of linkage mapping, QTL analysis, RNA-Seq data, and genome-wide association studies (GWAS) to explore relative flowering traits in almond (Prunus dulcis Mill. D.A. Webb). Horticultural Plant Journal.

67. Mrode, R. A., and Thompson, R. (2005). Linear models for the prediction of animal breeding values (2o).

68. Nelson, N. D., Berguson, W. E., McMahon, B. G., Cai, M., and Buchman, D. J. (2018). Growth performance and stability of hybrid poplar clones in simultaneous tests on six sites. Biomass and Bioenergy, 118, 115–125. 10.1016/j.biombioe.2018.08.007

69. Nsibi, M., Gouble, B., Bureau, S., Flutre, T., Sauvage, C., Audergon, J.-M., and Regnard, J.-L. (2020). Adoption and Optimization of Genomic Selection To Sustain Breeding for Apricot Fruit Quality. G3 Genes|Genomes|Genetics, 10(12), 4513–4529. 10.1534/g3.120.401452

70. Oakey, H., Verbyla, A., Pitchford, W., Cullis, B., and Kuchel, H. (2006). Joint modeling of additive and non-additive genetic line effects in single field trials. Theoretical and Applied Genetics, 113(5), 809–819. 10.1007/s00122-006-0333-z

71. Ortega, E., and Dicenta, F. (2003). Inheritance of self-compatibility in almond: Breeding strategies to assure self-compatibility in the progeny. Theoretical and Applied Genetics, 106(5), 904–911. 10.1007/s00122-002-1159-y

72. Ortega, E., and Dicenta, F. (2004). Suitability of Four Different Methods to Identify Self-Compatible Seedlings in an Almond Breeding Programme. The Journal of Horticultural Science and Biotechnology, 79(5), 747–753. 10.1080/14620316.2004.11511837

73. Pérez de los Cobos, F., Coindre, E., Dlalah, N., Quilot-Turion, B., Batlle, I., Arús, P., Eduardo, I., and Duval, H. (2023). Almond population genomics and non-additive GWAS reveal new insights into almond dissemination history and candidate genes for nut traits and blooming time. Horticulture Research, 10(10), uhad193. 10.1093/hr/uhad193

74. Pérez de los Cobos, F., Martínez-García, P. J., Romero, A., Miarnau, X., Eduardo, I., Howad, W., Mnejja, M., Dicenta, F., Socias i Company R., Rubio-Cabetas, M. J., Gradziel, T. M., Wirthensohn, M., Duval, H., Holland, D., Arús, P., Vargas, F. J., and Batlle, I. (2021). Pedigree analysis of 220 almond genotypes reveals two world mainstream breeding lines based on only three different cultivars. Horticulture Research, 8, 11. 10.1038/s41438-020-00444-4

75. Pérez de los Cobos, F., Romero, A., Lipan, L., Miarnau, X., Arús, P., Eduardo, I., Batlle, I., and Calle, A. (2024). QTL mapping of almond kernel quality traits in the F1 progeny of ‘Marcona’ × ‘Marinada’. Frontiers in Plant Science, 15. 10.3389/fpls.2024.1504198

76. Piaskowski, J., Hardner, C., Cai, L., Zhao, Y., Iezzoni, A., and Peace, C. (2018). Genomic heritability estimates in sweet cherry reveal non-additive genetic variance is relevant for industry-prioritized traits. BMC Genetics, 19(1), 23. 10.1186/s12863-018-0609-8

77. Piepho, H. P., Möhring, J., Melchinger, A. E., and Büchse, A. (2008). BLUP for phenotypic selection in plant breeding and variety testing. Euphytica, 161(1), 209–228. 10.1007/s10681-007-9449-8

78. Plummer, M., Best, N., Cowles, K., and Vines, K. (2006). CODA: convergence diagnosis and output analysis for MCMC. R news, 6(1), 7–11.

79. Poupon, V., Gezan, S. A., Schueler, S., and Lstibůrek, M. (2023). Genotype x environment interaction and climate sensitivity in growth and wood density of European larch. Forest Ecology and Management, 545, 121259. 10.1016/j.foreco.2023.121259

80. Prudencio, Á. S., Hoeberichts, F. A., Dicenta, F., Martínez-Gómez, P., and Sánchez-Pérez, R. (2021). Identification of early and late flowering time candidate genes in endodormant and ecodormant almond flower buds. Tree Physiology, 41(4), 589–605. 10.1093/treephys/tpaa151

81. Resende, M. D. V., Resende Jr, M. F. R., Sansaloni, C. P., Petroli, C. D., Missiaggia, A. A., Aguiar, A. M., Abad, J. M., Takahashi, E. K., Rosado, A. M., Faria, D. A., Pappas Jr., G. J., Kilian, A., and Grattapaglia, D. (2012). Genomic selection for growth and wood quality in Eucalyptus: Capturing the missing heritability and accelerating breeding for complex traits in forest trees. New Phytologist, 194(1), 116–128. 10.1111/j.1469-8137.2011.04038.x

82. Rutkoski, J. E. (2019). Estimation of Realized Rates of Genetic Gain and Indicators for Breeding Program Assessment. Crop Science, 59(3), 981–993. 10.2135/cropsci2018.09.0537

83. Sánchez-Pérez, R., Dicenta, F., and Martínez-Gómez, P. (2012). Inheritance of chilling and heat requirements for flowering in almond and QTL analysis. Tree Genetics & Genomes, 8(2), 379–389. 10.1007/s11295-011-0448-5

84. Sánchez-Pérez, R., Howad, W., Dicenta, F., Arús, P., and Martínez-Gómez, P. (2007b). Mapping major genes and quantitative trait loci controlling agronomic traits in almond. Plant Breeding, 126(3), 310–318. 10.1111/j.1439-0523.2007.01329.x

85. Sánchez-Pérez, R., Howad, W., Garcia-Mas, J., Arús, P., Martínez-Gómez, P., and Dicenta, F. (2010). Molecular markers for kernel bitterness in almond. Tree Genetics & Genomes, 6(2), 237–245. 10.1007/s11295-009-0244-7

86. Sánchez-Pérez, R., Ortega, E., Duval, H., Martínez-Gómez, P., and Dicenta, F. (2007a). Inheritance and relationships of important agronomic traits in almond. Euphytica, 155(3), 381–391. 10.1007/s10681-006-9339-5

87. Socias i Company R. (1998). Fruit tree genetics at a turning point: The almond example. Theoretical and Applied Genetics, 96(5), 588–601. 10.1007/s001220050777

88. Socias i Company R., and Felipe, A. J. (1992). Self-compatibility and autogamy in ‘Guara’ almond. Journal of Horticultural Science, 67(3), 313–317. 10.1080/00221589.1992.11516254

89. Socias, R., Company, I., Felipe, A. J., and Aparisi, J. G. (1999). A major gene for flowering time in almond. Plant Breeding, 118(5), 443–448. 10.1046/j.1439-0523.1999.00400.x

90. Sorensen, D. A., Andersen, S., Gianola, D., and Korsgaard, I. (1995). Bayesian inference in threshold models using Gibbs sampling. Genetics Selection Evolution, 27(3), 229–249.

91. Sorkheh, K., Shiran, B., Khodambashi, M., Moradi, H., Gradziel, T. M., and Martínez-Gómez, P. (2010). Correlations between quantitative tree and fruit almond traits and their implications for breeding. Scientia Horticulturae, 125(3), 323–331. 10.1016/j.scienta.2010.04.014

92. Tavassolian, I., Rabiei, G., Gregory, D., Mnejja, M., Wirthensohn, M. G., Hunt, P. W., Gibson, J. P., Ford, C. M., Sedgley, M., and Wu, S.-B. (2010). Construction of an almond linkage map in an Australian population Nonpareil × Lauranne. BMC Genomics, 11(1), 551. 10.1186/1471-2164-11-551

93. Vargas, F. J., Clavé, J., Romero, M., Batlle, I., and Rovira, M. (1998). AUTOGAMY STUDIES ON ALMOND PROGENIES. Acta Horticulturae, 470, 74–81. 10.17660/ActaHortic.1998.470.10

94. Vargas, F. J., and Romero, M. A. (1988). Comparación entre descendencias de cruzamientos intervarietales de almendro en relación con la época de floración y la calidad del fruto. 11557, 59–72.

95. Vargas, F. J., Romero, M. A., Rovira, M., and Girona, J. (1984). Amélioration de l’amandier par croisement de variétés. Résultats préliminaires à Tarragone (Espagne). 1, 101–122.

96. Vazquez, A. I., Bates, D. M., Rosa, G. J. M., Gianola, D., and Weigel, K. A. (2010). Technical note: An R package for fitting generalized linear mixed models in animal breeding. Journal of Animal Science, 88(2), 497–504. 10.2527/jas.2009-1952

97. Visscher, P. M., Hill, W. G., & Wray, N. R. (2008). Heritability in the genomics era—concepts and misconceptions. Nature Reviews Genetics, 9(4), 255–266

98. Ward, B. P., Brown-Guedira, G., Tyagi, P., Kolb, F. L., Van Sanford, D. A., Sneller, C. H., and Griffey, C. A. (2019). Multienvironment and Multitrait Genomic Selection Models in Unbalanced Early-Generation Wheat Yield Trials. Crop Science, 59(2), 491–507. 10.2135/cropsci2018.03.0189

99. Xu, S., and Xu, C. (2006). A multivariate model for ordinal trait analysis. Heredity, 97(6), 409–417. 10.1038/sj.hdy.6800885

100. Yang, H., and Su, G. (2016). Impact of phenotypic information of previous generations and depth of pedigree on estimates of genetic parameters and breeding values. Livestock Science, 187, 61–67. 10.1016/j.livsci.2016.03.001

