## SupplementalFile1 for "Dissecting the genetic architecture of flowering and maturity time in almond (*Prunus dulcis*): heritability estimates and breeding value predictions from historical data": SupplementalFile1.docx

**Initial Flowering Time (FI)**

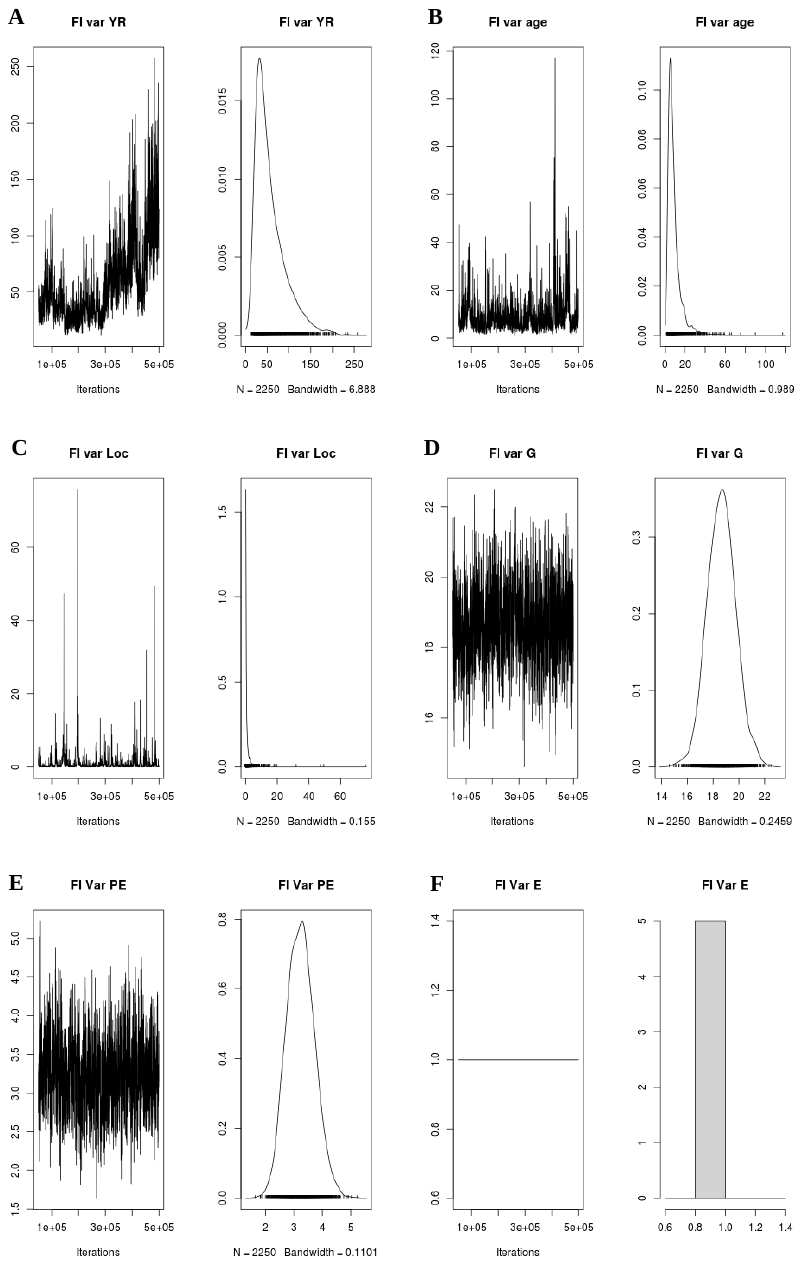

**Figure 1.** Markov chain trace plots and posterior density plots for the model effects of the Initial Flowering Time (FI) trait. **A.** Year effect (YEAR). **B.** Age effect (age). **C.** Parcel or location effect (Loc). **D.** Additive genetic effect (G). **E.** Permanent environmental effect. **F.** Residual effect (e). The symmetrical and non-aberrant curves indicate good chain convergence and unimodal posterior distributions.

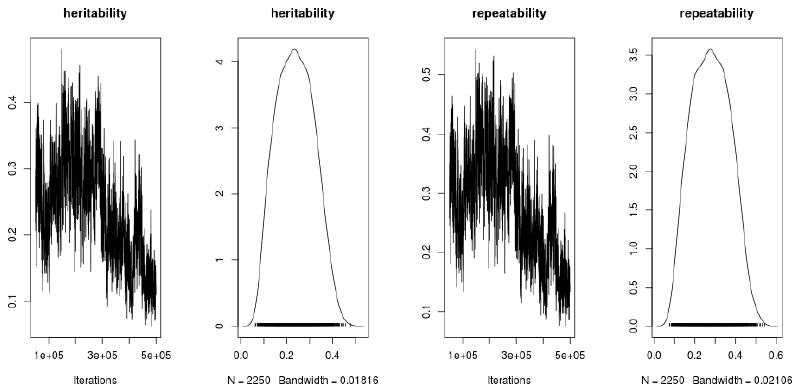

**Figure 2.** Markov chain trace plots and posterior density plots for the heritability and repeatability parameters of the Initial Flowering Time (FI) trait. The symmetrical and non-aberrant curves indicate good chain convergence and unimodal posterior distributions.

**Full Flowering Time (FP)**

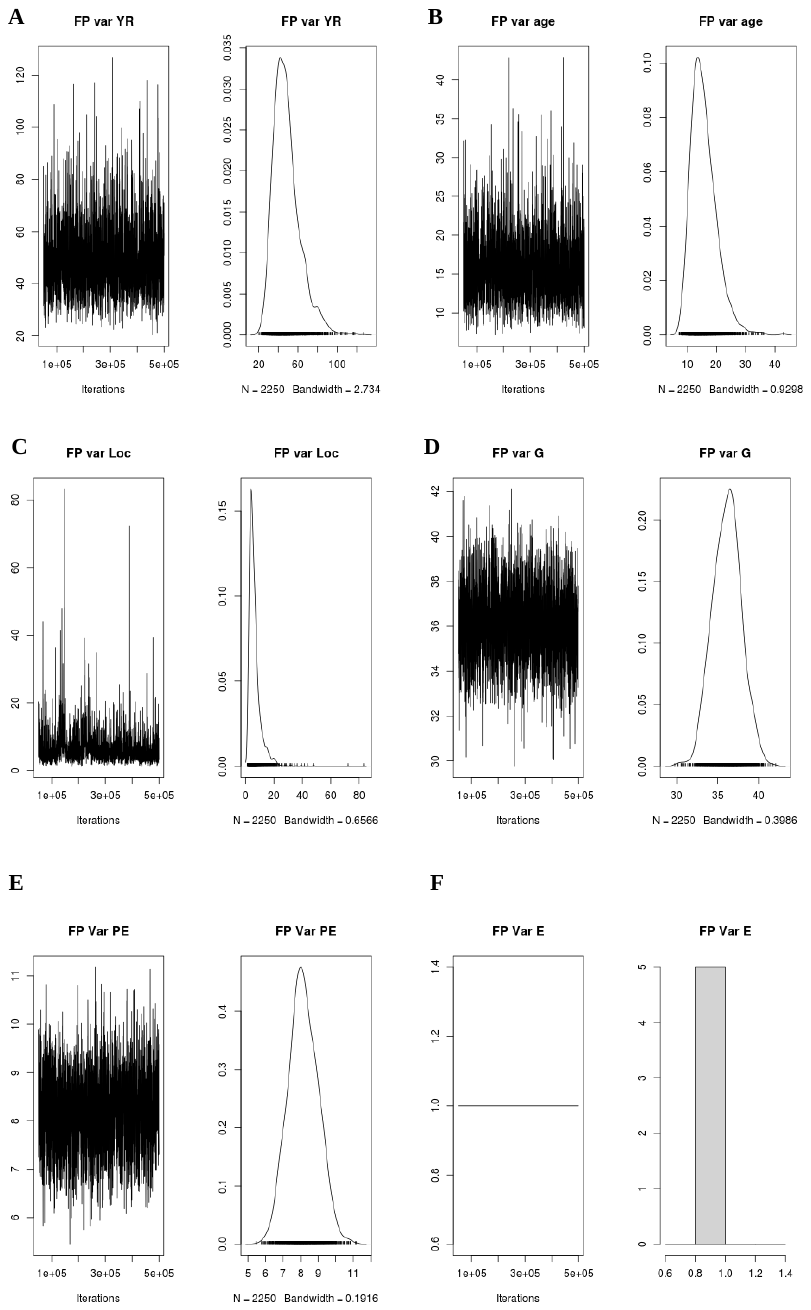

**Figure 3.** Markov chain trace plots and posterior density plots for the model effects of the Full Flowering Time (FP) trait. **A.** Year effect (YEAR). **B.** Age effect (age). **C.** Parcel or location effect (Loc). **D.** Additive genetic effect (G). **E.** Permanent environmental effect. **F.** Residual effect (e). The symmetrical and non-aberrant curves indicate good chain convergence and unimodal posterior distributions.

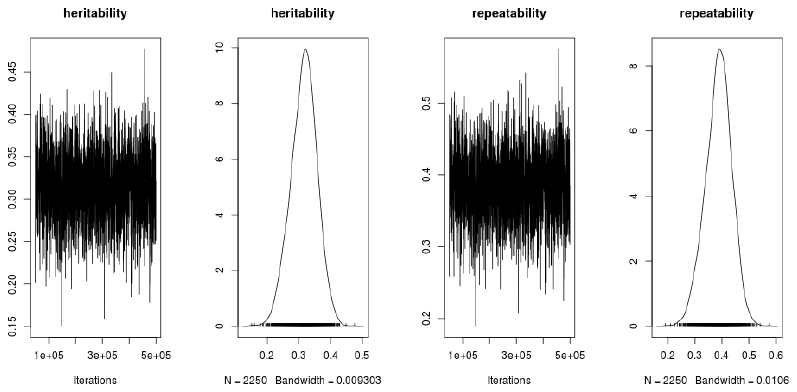

**Figure 4.** Markov chain trace plots and posterior density plots for the heritability and repeatability parameters of the Full Flowering Time (FP) trait. The symmetrical and non-aberrant curves indicate good chain convergence and unimodal posterior distributions.

**Final Flowering Time (FF)**

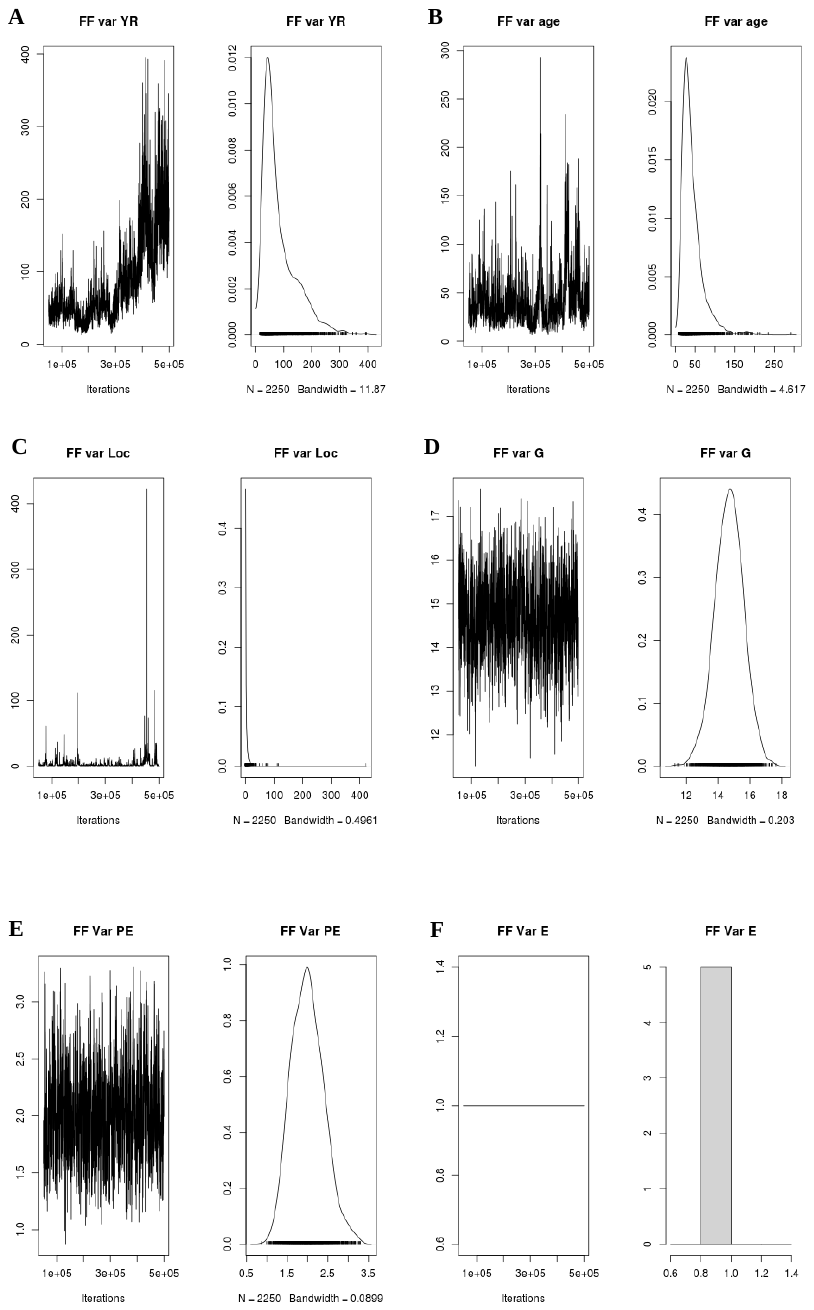

**Figure 5.** Markov chain trace plots and posterior density plots for the model effects of the Final Flowering Time (FF) trait. **A.** Year effect (YEAR). **B.** Age effect (age). **C.** Parcel or location effect (Loc). **D.** Additive genetic effect (G). **E.** Permanent environmental effect. **F.** Residual effect (e). The symmetrical and non-aberrant curves indicate good chain convergence and unimodal posterior distributions.

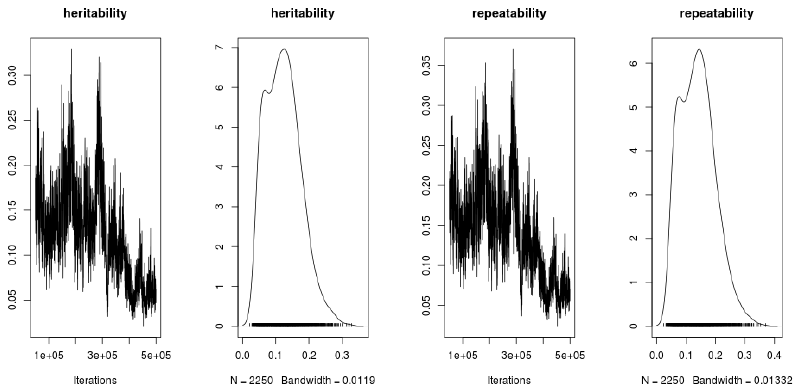

**Figure 6.** Markov chain trace plots and posterior density plots for the heritability and repeatability parameters of the Final Flowering Time (FF) trait. The symmetrical and non-aberrant curves indicate good chain convergence and unimodal posterior distributions.

**Maturity Time (MADUR)**

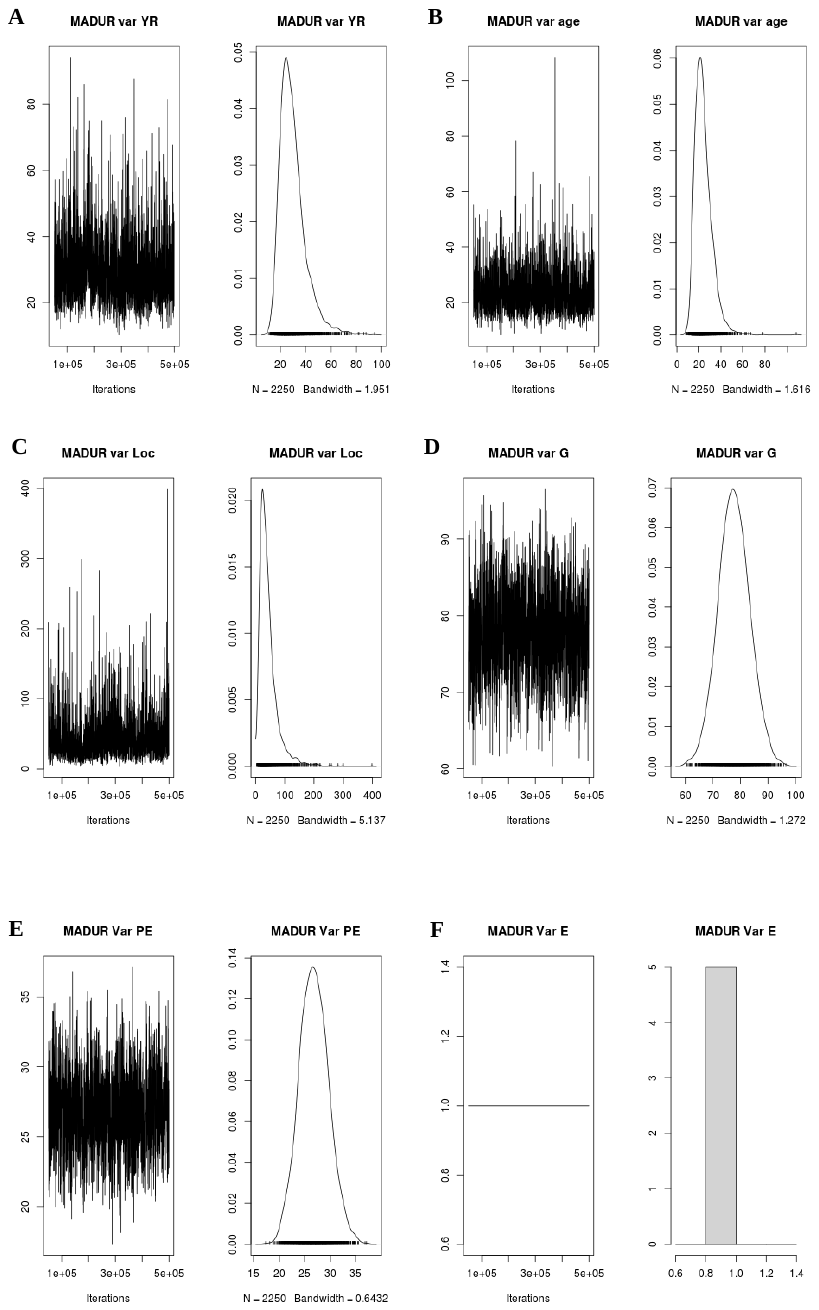

**Figure 7.** Markov chain trace plots and posterior density plots for the model effects of the Maturity Time (MADUR) trait. **A.** Year effect (YEAR). **B.** Age effect (age). **C.** Parcel or location effect (Loc). **D.** Additive genetic effect (G). **E.** Permanent environmental effect. **F.** Residual effect (e). The symmetrical and non-aberrant curves indicate good chain convergence and unimodal posterior distributions.

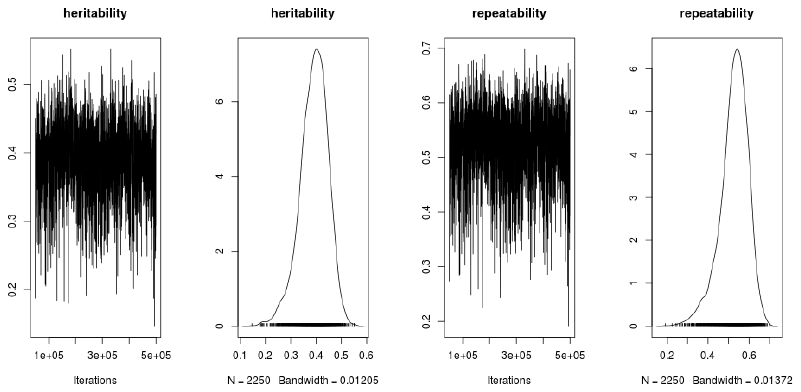

**Figure 8.** Markov chain trace plots and posterior density plots for the heritability and repeatability parameters of the Maturity Time (MADUR) trait. The symmetrical and non-aberrant curves indicate good chain convergence and unimodal posterior distributions.

**Flower Density (FLINT)**

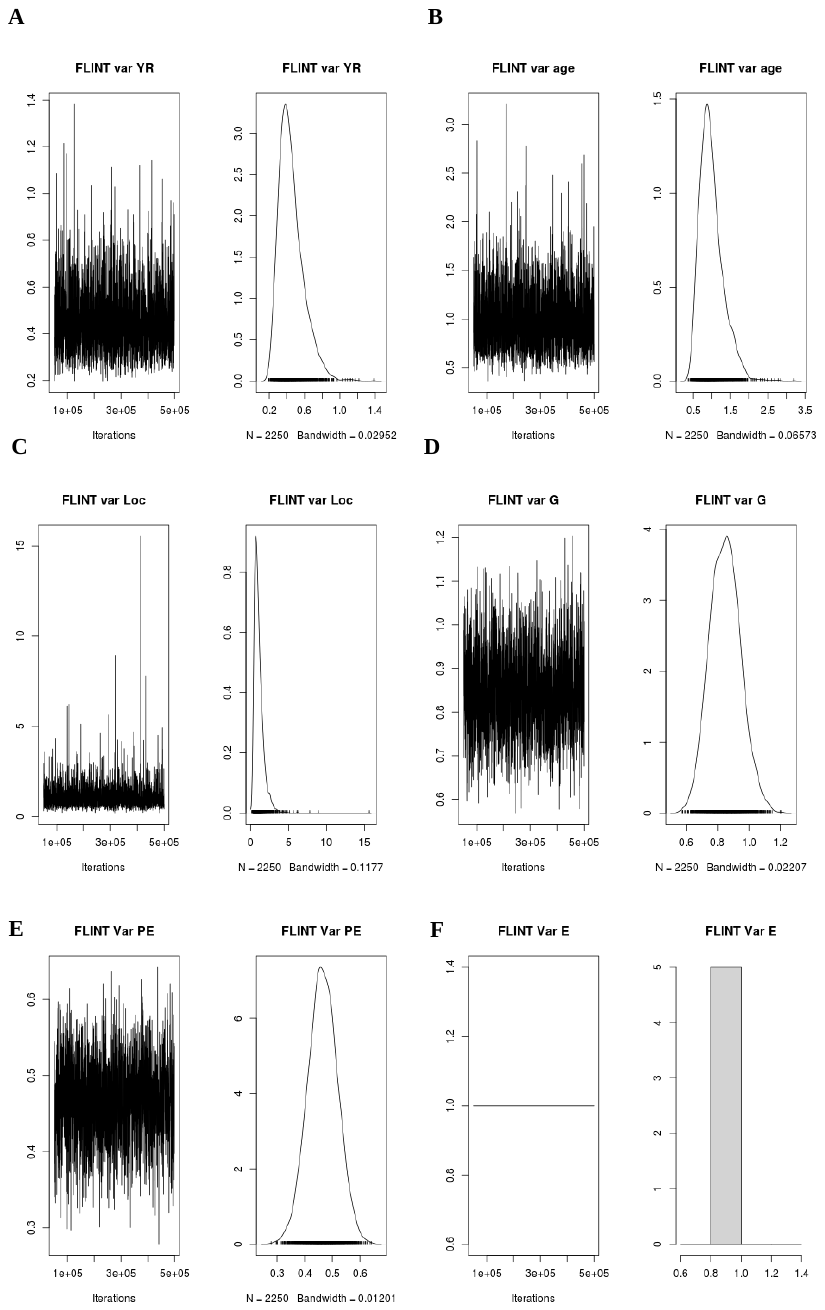

**Figure 9.** Markov chain trace plots and posterior density plots for the model effects of the Flower Density (FLINT) trait. **A.** Year effect (YEAR). **B.** Age effect (age). **C.** Parcel or location effect (Loc). **D.** Additive genetic effect (G). **E.** Permanent environmental effect. **F.** Residual effect (e). The symmetrical and non-aberrant curves indicate good chain convergence and unimodal posterior distributions.

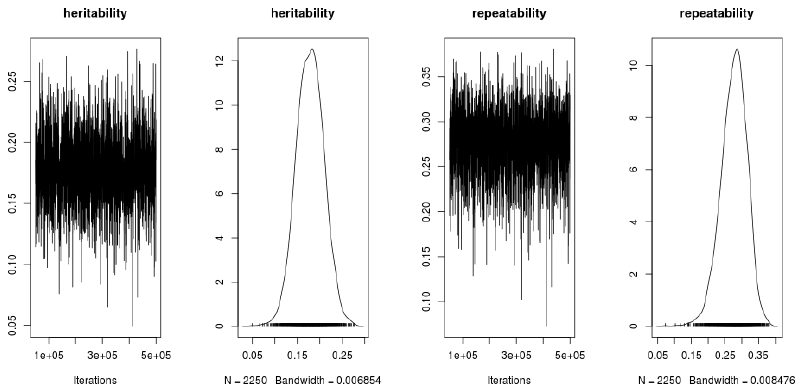

**Figure 10.** Markov chain trace plots and posterior density plots for the heritability and repeatability parameters of the Flower Density (FLINT) trait. The symmetrical and non-aberrant curves indicate good chain convergence and unimodal posterior distributions.
